# Prediction variability in physiologically based pharmacokinetic modeling of tissue disposition under deep uncertainty

**DOI:** 10.64898/2025.12.05.692437

**Authors:** Mustafa Farahat, DT Flaherty, Zachary R Fox, Belinda S Akpa

## Abstract

Physiologically based pharmacokinetic (PBPK) models are increasingly invoked in virtual screening workflows for therapeutics. These mechanistic models project pharmacokinetic outcomes from molecular properties, with data-driven models acting as intermediaries to map molecular structure to PBPK input parameters. Errors in predicted parameters and unvalidated assumptions within PBPK models expose PK predictions to deep uncertainty. Herein, we examine how these uncertainties affect the prediction variability of dynamic, tissue-specific exposure. We validated four PBPK models against 1,854 experimental datapoints – to establish their predictive fidelity before introducing parameter uncertainty typical of property-prediction models. Depending on molecule properties and model choice, the coefficient of variation under parameter uncertainty ranged from 10^-6^ to 31 for predicted PK statistics. Further, we identified notable model disagreement for a subset of drug-like chemical space characterized by lipophilic, protonated molecules. Uncertainty quantification revealed biophysicochemical properties and parameter interactions that drove disagreement and highlighted model assumptions that exacerbated prediction variance. Our findings delineate the challenges presented by deep epistemic uncertainty in PBPK modeling.

## Introduction

AI-driven drug discovery is evolving rapidly as academic groups, industry, and start-ups develop computational tools to generate and evaluate candidate molecules^1–4^. Computational approaches can shift screening paradigms from sequential experimentation to parallel exploration^5–7^, making it possible to evaluate candidate structures through workflows that integratively account for molecular activity^8,9^, identify off-target interactions^10–12^, and discriminate toxicity liabilities^13,14^ – amongst other therapeutic criteria. Predicting these properties directly from chemical structures is a key step that supports high-throughput screening before synthesis, empirical development, and *in vivo* testing^15,16^. However, whether such predictions accelerate discovery depends on the ability of constituent models to capture the features that make a molecule a successful therapeutic. For a molecule to function as a drug, it must engage in desired molecular interactions, reach its site of action within the body, and persist there at concentrations sufficient to modulate biological processes^17–19^. Many efforts in the domain of AI-driven discovery focus on data-driven^20,21^ and surrogate models^22,23^ that predict molecular-scale phenomena governing candidate potency and safety. What may be neglected in these workflows are the dynamic features of human physiology that determine whether a compound can reach its target *in vivo*, at high enough concentrations, and for long enough to influence disease processes^17,24,25^. Physiologically based pharmacokinetic (PBPK) modeling can address these issues by mapping molecular properties to chemical fate in the human body.

PBPK models are systems of ordinary differential equations that mechanistically describe the body as an assembly of organ compartments connected by circulating blood^26–30^. Each equation takes the form of a material balance capturing the influence of tissue-specific composition on molecule distribution and metabolism. Solving the system of equations thus yields tissue-specific, time-resolved predictions of drug exposure – pharmacokinetic outcomes directly relevant to therapeutic efficacy. As a well-established tool in later stages of drug development^28,31–33^, PBPK modeling supports *post hoc* rationalization of therapeutic outcomes and helps refine therapeutic regimens^34,35^. The models may be used to prospectively explore clinical dosing strategies, assess drug-drug interactions, or predict outcomes for special populations such as pediatric patients or those with impaired organ function^36–39^. Their informational value in these late-stage applications has driven growing efforts to deploy PBPK models in virtual screening. Emerging screening workflows now combine machine learning (ML) models that predict molecular properties with PBPK frameworks to project *in vivo* drug behavior directly from chemical structures^40–46^, and possibly augment the decision-making value of novel alternative methods^47–50^. However, applying PBPK models in this way introduces new challenges. When applied at the clinical stage or later, these models rely on experimentally measured parameters and can be validated against drug– and species-specific observations. When PBPK models are instead used in ML-informed workflows, establishing their credibility becomes more difficult. A primary issue is the impact of uncertainty arising from two sources.

The first source is structural. Any PBPK model is an abstraction of the true physiological system^51^. When applied broadly across diverse chemical space, such assumptions may misrepresent key physiological processes in ways that are not easily detected without experimental validation^26,52,53^. For practical decision-making in virtual screens, a single model – or small set of models – would need to produce informative predictions across a wide range of chemical structures. The second source of uncertainty is parametric. For virtual compounds, drug properties cannot be determined experimentally and instead drug-specific input parameters for PBPK modeling can be obtained from data-driven quantitative structure-activity relationship (QSAR) models^54,55^. These approaches are often trained on sparse or biased datasets and can lead to large, compound-specific prediction errors. As a result, both the PBPK framework and its input parameters produce a scenario of deep epistemic uncertainty. Understanding how this uncertainty propagates through PBPK frameworks is key to judging whether model predictions are useful in early virtual-screening workflows.

In this study, we assess the confidence associated with PBPK predictions for virtual compounds, where both model assumptions and input parameters are uncertain. To this end, we examined: model prediction fidelity (for known drugs with empirically characterized input parameters) and prediction variance (for virtual compounds whose input parameters are perturbed to reflect the uncertainty typical of published QSAR models). We quantified prediction fidelity by evaluating each model’s ability to predict plasma-tissue partitioning coefficients (*Kp*) and volume of distribution (*Vss*) – both equilibrium PK parameters with available empirical data for model validation. Then, with ML-informed PBPK workflows in mind, we introduced parameter uncertainty representative of the mean absolute error reported in published property-prediction models^56–58^. We propagated this uncertainty through four PBPK models: (1) our own model, modified from the Rodgers & Rowland (R&R)-Lukacova framework, (2) a second R&R-Lukacova based model reported by Mathew *et al.*^59^, and (3 & 4) the calibrated and uncalibrated variants of the Schmitt-based model reported by Pearce *et al.*^60^. Using Monte Carlo simulations, we quantified tissue-specific PK statistics relevant to intracellular drug exposure for a synthetic dataset of 10^4^ virtual molecules. Specifically, for each molecule, we recorded steady-state volume distribution (*Vss*), maximum unbound concentration (*Cmax,u*), area under the unbound concentration-time curve (*AUCu_0-∞_*), and the duration of time for which unbound concentration exceeded 0.1 μM (*T_0.1μM_*). We thereby quantified the impact of epistemic uncertainty on prediction variance of model-derived PK outcomes.

Our analysis of prediction variance addressed three core questions: (1) Do PBPK models embedding different mathematical abstractions for tissue partitioning exhibit different susceptibility to parameter uncertainty? (2) If so, where in chemical space do these predictions differ most, and is that divergence evident for nominal predictions or specifically induced under parameter uncertainty? (3) Which drug-specific input parameters contribute most to prediction variability and model disagreement, and where would efforts to improve property prediction models be most impactful?

To answer these questions, we (1) evaluated model agreement, (2) identified regions of chemical space with divergent predictions, and (3) applied global sensitivity analysis to quantify the contribution of each input parameter to prediction uncertainty. We have thereby shed light on challenges to consider when using ML-informed PBPK workflows to guide compound assessment, particularly when parameter uncertainty is high and empirical validation is not feasible.

## Results and Discussion

Mechanistic models have been developed to predict one critical class of drug-specific PBPK parameters: plasma-tissue partition coefficients (*Kp*). These constants describe the equilibrium partitioning of drug between the blood plasma and a given tissue and can be predicted as a function of tissue composition. Partitioning models may have an explicitly or implicitly bounded domain of applicability or contain terms that must be selected based on molecular properties. Some work well for acids and zwitterions, others for strong bases, and still others for highly lipophilic compounds^61–64^. Most are designed to address known drug-like compounds, while others aim to be predictive over a broader range of molecules from drugs to chemicals of concern for environmental exposure^65^. In all cases, prediction of *Kp* requires biophysicochemical property inputs for the molecule of interest. For PBPK models that are used early in the discovery pipeline, QSAR models provide a data-driven mapping of molecule structure to molecular properties.

In this study, we interrogate PBPK models based on three tissue-partitioning models: 2 canonical approaches and a modified variant introduced herein. As reference approaches, we consider the model of Mathew *et al.*^59^ and that of Pearce *et al.*^60^. These represent, respectively, modifications of the R&R-Lukacova framework and the Schmitt framework for predicting tissue-plasma partitioning. The model we introduce herein is based on the R&R-Lukacova framework, adopting alternative assumptions for drug association with proteins (interstitial and intracellular), neutral lipids, and phospholipids. The distinguishing features of the three *Kp* models are presented in the Methods section (**Table 1**). In considering a modified formulation for tissue partitioning relationships, we aimed to improve the model’s ability to generate predictions of similar quality across drug-like chemical space.

**Table 1.** Distinguishing features of the models considered in this work. *MA*: membrane affinity. *Dow*: octanol-water distribution coefficient. *logD*: distribution coefficient. *lopP*: partition coefficient. AP: acidic phospholipid. R&R: Rogers & Rowland. *fu*: fraction unbound.

|  | This model | Pearce (2017) | Mathew (2021) |
| --- | --- | --- | --- |
| **fu adjustment** | Modified Lukacova equation to use $D_{ow}$ for neutral lipid partitioning, membrane affinity (MA) for neutral phospholipids, and to include plasma acidic phospholipids. | Addresses lipid partitioning in plasma using an effective volume fraction of neutral lipid that includes 30% of the neutral phospholipid volume. | Weights lipid partitioning by the fraction of molecules in a neutral state. Adjusts protein binding term to reflect neutral and charged fractions binding to plasma lipoprotein and albumin, respectively. |
| **intracellular pH** | Accounts for pH differences in intracellular compartments of different tissues. | Accounts for pH differences in intracellular compartments of different tissues. | pH=7 in intracellular compartment of all tissues. |
| **neutral lipid partitioning** | Describes partitioning into neutral lipids using $logD$ at relevant pH. | Describes partitioning into neutral lipids using $logD$ at relevant pH. | Assumes only neutral drugs partition into neutral lipids with partitioning described by $logP$. |
| **neutral phospholipid partitioning** | Partitioning into neutral phospholipids described using the $logMA$ (membrane affinity) correlation of Yun and Edginton. Employs $logD$ in this correlation, to reflect impact of drug ionization. | Partitioning into neutral phospholipids described using the $logMA$ (membrane affinity) correlation of Yun and Edginton. Employs $logP$ in this correlation. | Only neutral drugs partition into neutral phospholipids, with partition coefficient dependent on $logP$ (*viz.* $0.3P+0.7$). |
| **acidic phospholipid partitioning** | Assumes all molecules associate with acidic phospholipids, with coefficient dependent on ionization, as described by Schmitt. Zwitterionic fractions grouped with the cationic fraction in AP partitioning (following the assumption of R&R, which treats protonated sites as dominating AP interactions for zwitterions). | Assumes all molecules associate with acidic phospholipids, with coefficient dependent on ionization, as described by Schmitt. Zwitterionic fraction treated as part of the neutral fraction in AP partitioning. | Assumes drug fractions with cationic sites bind to acidic phospholipids, regardless of overall molecular charge. |
| **interstitial protein binding** | Assumes anionic and zwitterionic fractions bind to albumin and neutral drug fractions bind to lipoproteins. Derives corresponding protein binding affinity from $f_u$. Interstitial protein content varies with tissue albumin and lipoprotein levels. | Assumes nonspecific binding to proteins. Derives corresponding protein binding affinity from $f_u$. Interstitial protein content is invariant across tissues. | Assumes all ionized drug fractions bind to albumin and neutral fractions bind to lipoproteins. Derives protein binding affinity from $f_u$. |
| **intracellular protein binding** | Assumes nonspecific binding to proteins using an empirical relationship linking protein binding to phospholipid partitioning. | Assumes nonspecific binding to proteins using an empirical relationship linking protein binding to phospholipid partitioning. | No intracellular protein binding, except in blood cells. In blood cells, non-cations assumed to bind to an unspecified protein, with affinity equal to that of cation binding to acidic phospholipids. |
| **adjusted lipid partitioning for adipose** | Replaces octanol-water partition coefficient with the vegetable oil-water partition coefficient. | N/A | Replaces octanol-water partition coefficient with the vegetable oil-water partition coefficient. |

### Model fidelity-to-data

To assess model performance, we quantified four common statistics that capture fidelity with respect to experimentally determined equilibrium PK quantities *–* namely, tissue partitioning coefficients and volume of distribution. We subsequently quantified each model’s: (1) *accuracy*, as indicated by the fraction of drugs with PK predictions within 2-fold, 3-fold, and 10-fold error; (2) *precision*, as indicated by the root mean squared error of log-transformed values, RMSLE; (3) *bias*, as indicated by average fold error, AFE; and (4) *agreement*, as indicated by the concordance correlation coefficient, CCC. CCC describes the extent to which predictions fall on the identity line of a parity plot (*i.e.*, when predictions are plotted against measured quantities). As with other correlation coefficients, CCC ranges between –1 and 1. We provide equations for each of these performance metrics in the Methods section.

#### Data sources

PBPK models have three classes of parameter: (1) those that reflect the nature of the biological system (*e.g.*, tissue volumes and composition), (2) those that solely reflect the physicochemical properties of the drug (*e.g.*, distribution coefficient, *logD*, and acid dissociation constants, *pKa*), and (3) those that capture interactions between drug chemistry and biology (*e.g.*, metabolic clearance, *CL_int_*, and plasma protein binding, *fu*). **Table S1** summarizes each dataset used herein, its source, and its application within this study.

### System-specific data

System-specific parameters for rat and human physiology were taken from the Httk *physiology.data* table^66^. Likewise, tissue composition information was adapted from Httk (*tissuedata.data*)^66^. Data describing tissue vascular fraction and tissue albumin and lipoprotein concentrations were drawn from Edginton *et al*. and Rodgers *et al.*, respectively^67,62^

### Drug-specific data

To assess the fidelity of *Kp* predictions, we sourced rat tissue partitioning data from the dataset collated by Pearce *et al.*^60^. This dataset includes 157 distinct compounds over 11 tissues, with a total of N = 992 data points. For *Vss*, we used the human *in vivo* values reported by Obach *et al.*^68^ *–* limiting our predictions to the 862 compounds for which those authors present the required set of drug-specific input parameters. **Figure S1** shows the distributions of these molecular properties in the Pearce and Obach datasets.

#### Model calibration

A key feature of the Pearce model is the use of an empirical correction to adjust predicted values of *Kp*^60^. These corrections were derived from linear regressions of predicted versus measured tissue-partition coefficients for 157 pharmaceutical compounds across 11 tissues. The resulting tissue-specific linear corrections yield a calibrated set of *Kp* values that are intended to improve overall prediction fidelity.

To assess this model’s fidelity on equal footing with the other *Kp* models (*i.e.*, those that have not been “tuned” to that empirical dataset), we evaluated its performance based on the uncalibrated *Kp* predictions. We then used the calibrated form of the Pearce model when assessing the credibility of human *Vss* predictions.

#### Canonical models exhibit similar fidelity to rat *Kp* and human *Vss* data

When tallied over all 157 drugs and 11 tissues in the *Kp* dataset, the Mathew model underperformed in predicting rat tissue partitioning coefficients (**Figure 1A**). Less than 40% of *Kp* values were predicted within a 2-fold error. The Mathew model also failed to exceed the 90% mark for predictions within a 10-fold error. Although our model shares an underlying framework with Mathew (*i.e.*, that of R&R-Lukacova), it performed with accuracy comparable to that of uncalibrated Pearce. When we disaggregated the drug molecules by ion class, it became clear that the overall performance of the Mathew model was disproportionately impacted by poor predictions for zwitterions – for which less than 20% were predicted within a 2-fold error. Across acids, bases, and neutral compounds, we observed more parity in performance among the three models. However, the Mathew model did not yield greater accuracy than our model or Pearce’s for any of the ion classes. The detailed performance statistics for rat *Kp* across ion classes are presented in **Table S2**.

**Figure 1.**
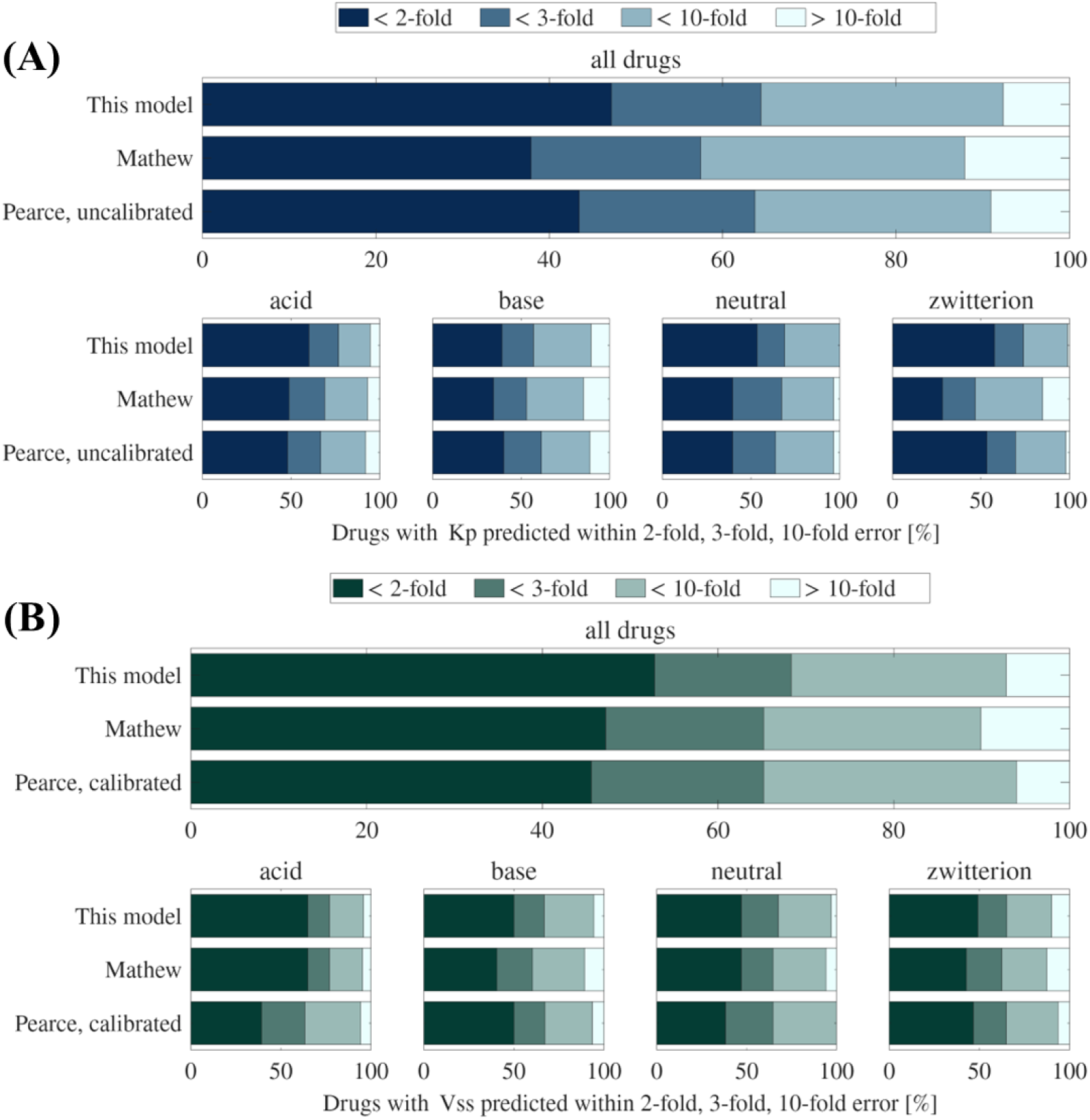
Fold error statistics for each model and ion class. (A) Rat *Kp*. Statistics summarize the quality of 992 predictions per model, spanning 157 molecules and 11 tissues. (B) Human *Vss*. Statistics summarize the quality of 862 predictions per model – one *Vss* prediction per molecule.

Note that these fold-error quantities should be interpreted in the context of the number of experimental measurements reported for compounds in each ion class. By far the largest group was bases (N=584), followed by acids (N=258), zwitterions (N=92), and neutrals (N=58). At the two-fold error level our model appears to perform comparatively well on neutral compounds. However, the ∼14% difference between our model and Pearce’s uncalibrated model corresponds to only 8 data points. As the validation data does not provide a measurement for every tissue across the full set of drugs, this difference of 8 data points could represent 8 different compounds, with empirical measurements within the same tissue, or the same compound measured over 8 different tissues (or, indeed, distinct compounds across distinct tissues). Thus, we interpreted the relative accuracy of the *Kp* predictions with caution.

When we inspected human *Vss* predictions, we again noted similar accuracy between our model and that of Pearce (now the calibrated model), with the Mathew model lagging in performance (**Figure 1B**). When disaggregated, however, the Mathew model maintained something closer to parity across all ion classes. There is comparatively poorer performance again for zwitterions, but it is less marked than in the *Kp* evaluation. For *Vss*, the underperformance with respect to zwitterions is likely dampened by the fact that this PK metric is a composite quantity, computed as a *Kp*-weighted sum over the volumes of individual tissues (**Equation 19**). Tissues for which equilibrium partitioning constants are poor reflections of relevant biological transport phenomena^69–72^ (*e.g.*, brain, liver) represent relatively small contributions by volume^66,73^. By contrast, tissues that represent disproportionately large volumes (*e.g.*, muscle, adipose) are relatively well described by equilibrium partitioning^74,75^. The full set of *Vss* performance statistics for each ion class is summarized in **Table S3**.

In **Figure 2**, we compare disaggregated *Kp* predictions for the brain, adipose, and muscle compartments. These tissues represent ∼2%, ∼24%, and ∼40% of the body’s volume, respectively^66,73^. As expected, the predictive accuracy for brain tissue was relatively poor, with CCC below 0.6 in all models. Correlations for adipose *Kp* predictions were the best out of all tissues, for all models. This is despite the models invoking different approaches to describe neutral lipid partitioning in adipose tissue. Our model and that of Mathew follow the practice of adopting the vegetable-oil-water partition coefficient to capture the high triglyceride content of that tissue. The Pearce model does not make this adjustment. The Pearce model does, however, use *logD* rather than *logP* to describe lipid partitioning – a feature that is shared by our model. Mathew differs in that *logP* is retained, but the partitioning is only associated with the fraction of molecules predicted to be in the uncharged state.

**Figure 2.**
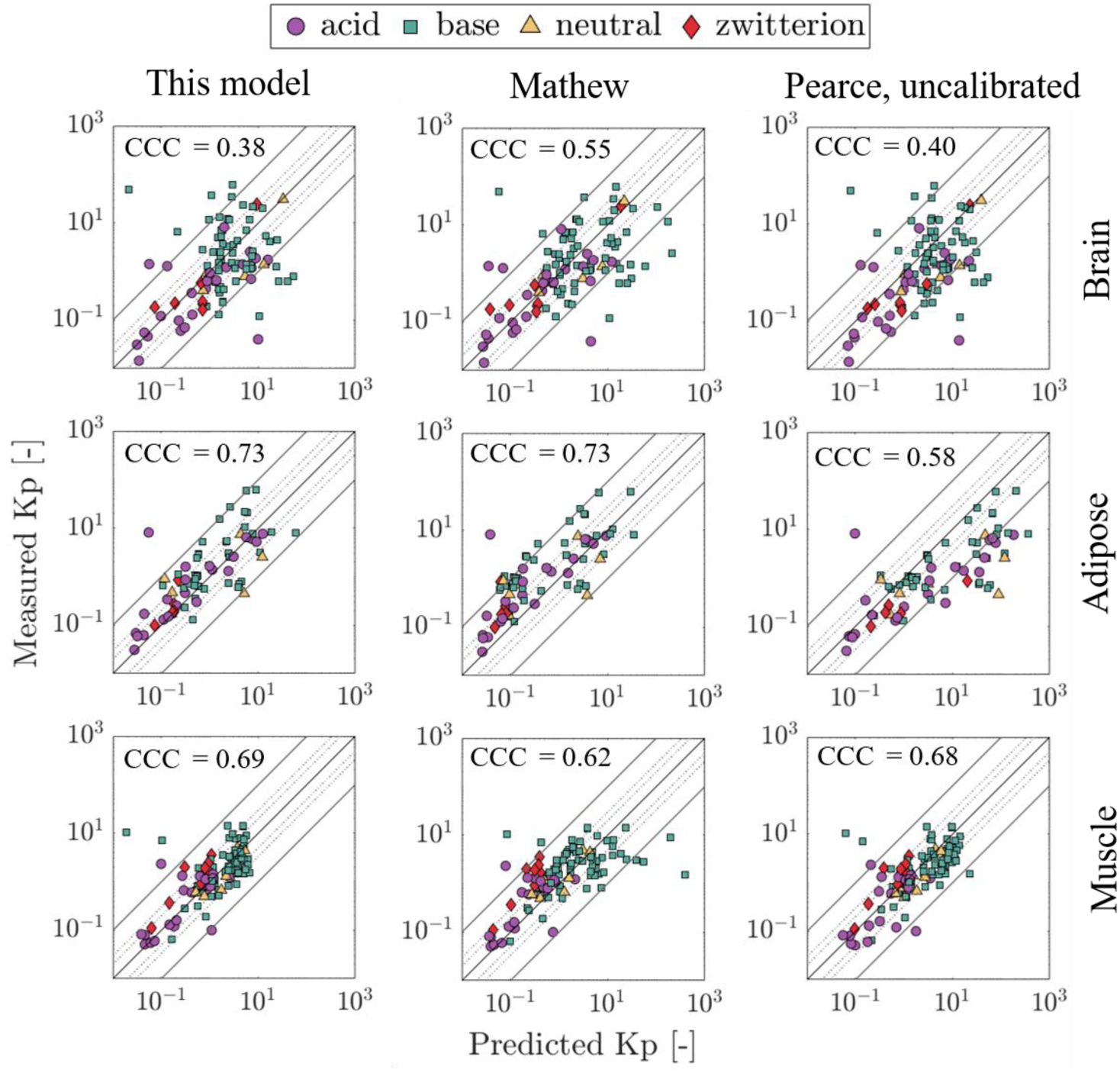
Measured vs. predicted rat tissue partitioning coefficients (*Kp*). Left: This model. Center: Mathew model. Right: Uncalibrated Pearce model. Lines indicate the identity line, two-fold, three-fold, and ten-fold error intervals.

Figure 3A summarizes the RMSLE, AFE, and CCC performance statistics for the complete set of *Kp* predictions. Across all ion classes except bases, our model exhibits the lowest RMSLE (*i.e.*, highest precision), along with a consistent tendency to underpredict tissue partitioning (*i.e.*, log_2_AFE < 0). This tendency is shared by the other R&R-Lukacova model, *i.e.*, that of Mathew, which yields the greatest prediction biases. Overall *Kp* prediction bias is least pronounced for the Pearce (uncalibrated) model. This pattern is not replicated across the *Vss* predictions (Figure 3B). Here, for all ion classes, prediction bias is greatest for the calibrated Pearce model, where *Vss* tends to be overpredicted.

**Figure 3.**
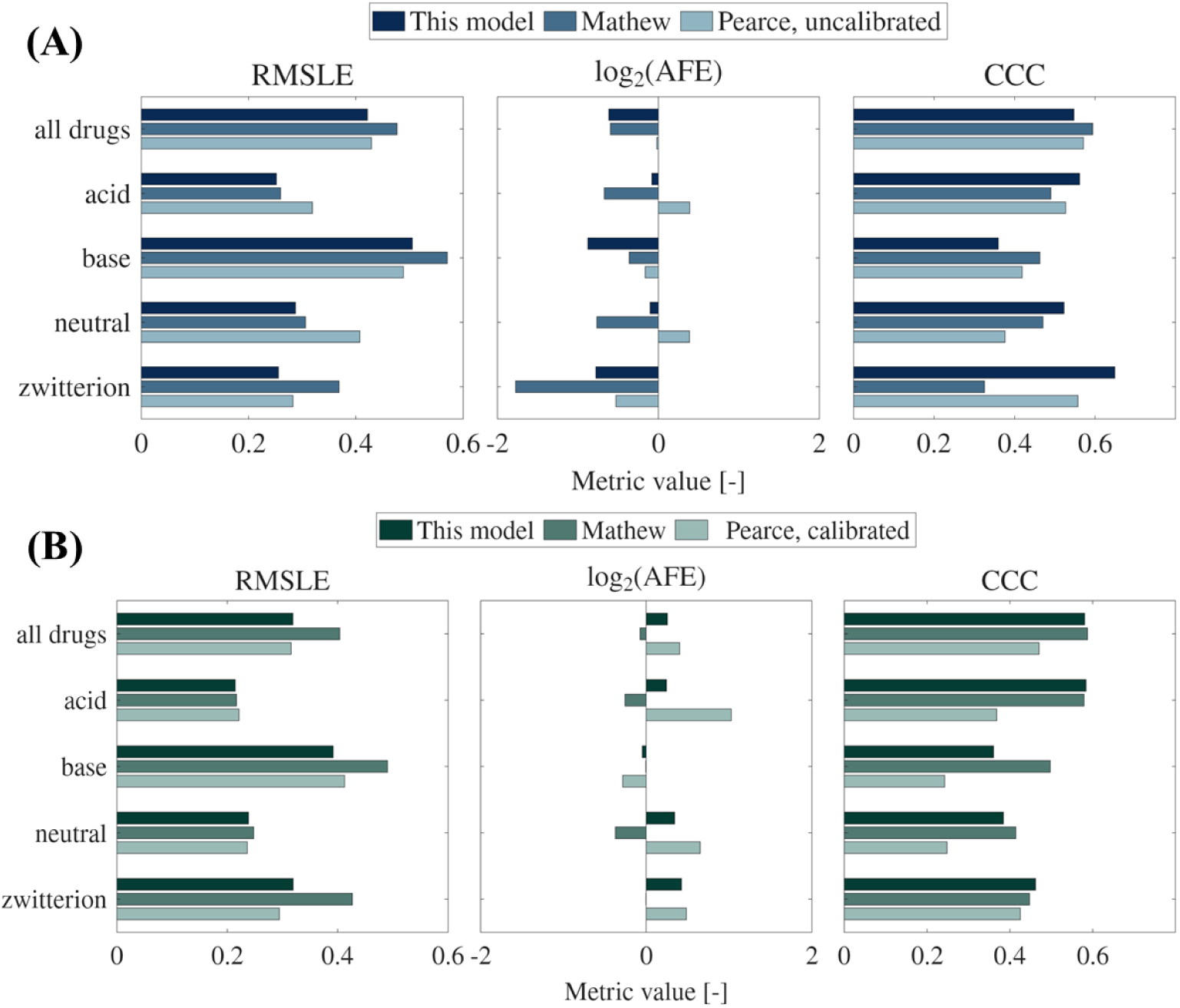
Precision, bias, and correlation statistics for each model and ion class. (A) Rat *Kp*. Statistics summarize the quality of 992 predictions per model, spanning 157 molecules and 11 tissues. (B) Human *Vss*. Statistics summarize the quality of 862 predictions per model – one *Vss* prediction per molecule.

In short, despite the non-trivial differences in underlying framework and assumptions, all three models yield predictions of broadly similar fidelity to data. Differences are, however, observed at the level of individual tissues. These tissue-level errors are obscured at the level of *Vss*, as *Kp* predictions for the largest two tissues have high fidelity. Tissue-specific errors will, however, be of concern for evaluating any tissue-specific exposure criterion. Further differences emerge when results are disaggregated by ionization class. Our analysis demonstrates that specific assumptions modulate error and bias within a PBPK framework and the choice of underlying framework from amongst canonical approaches is not a defining determinant of model credibility. This conclusion, however, assumes perfect knowledge of each drug’s properties. Within an ML-informed PBPK workflow, drug-specific input parameters (*e.g.*, RMM, *fu*, *CL_int_*, *logD*, donor and acceptor *pKa*) are obtained from QSAR models that map chemical structure to the molecular properties required for PBPK simulation. Data reflecting human biology can be limited and highly variable, so data-driven prediction of parameters *fu* and *CL_int_* proves particularly recalcitrant^76–81^. Identifying informative molecular features and predictive modeling strategies to generate these drug-specific parameters is a foundational requirement for reliable predictions – and thus an active area of research^82–85^. In the following sections, we examine the impact that prediction errors typical of existing property-prediction models would have on the uncertainty of PBPK predictions.

### Prediction variance under epistemic uncertainty

To be useful as a tool for decision making in an ML-informed PBPK workflow, a PBPK model would need to exhibit two key properties: (1) the model should be adequately sensitive to variations in molecular properties to reflect the impact of those variations on PK outcomes of therapeutic consequence, and (2) despite uncertainty in predicted parameter values, those PK predictions should remain informative about the relative PK performance of virtual candidates. The prior section addressed the first concern by evaluating PBPK model fidelity for known compounds with measured biophysicochemical parameters – considering the epistemic uncertainty associated with model formulation. We now turn to the second concern by examining how PBPK predictions respond to uncertainty in their input parameters. Specifically, we assess whether predictions retain their informative value once uncertainty is introduced and whether parameter uncertainty has disparate impacts on model predictions in particular regions of chemical space.

To do so, we generated 10^4^ pseudomolecules by correlated random sampling of *RMM*, *fu, CL_int_, logD,* and *pKa* from property distributions characterizing the compounds in the Obach dataset (see pseudocode in **Algorithm 1**). We then evaluated the impact of parameter uncertainty by performing Monte Carlo simulations using perturbed molecular properties. To this end, for each pseudomolecule, we assumed a parametric distribution centered on the molecule’s “true” property value and scaled its spread to match the mean absolute error (MAE) reported for published QSAR models^56–58^ (see **Table 2**). *LogD* was sampled from a normal distribution, *fu* and *pKa* from beta distributions, and *CL_int_* from a lognormal distribution. We ran 10^3^ Monte Carlo realizations per pseudomolecule through each of the four PBPK models. The resulting unbound concentration-time profiles allowed us to compute outcome distributions for *Vss* and dynamic tissue-specific PK metrics *Cmax,u*, *AUCu*, and *T_0.1μM_*.

**Table 2.**
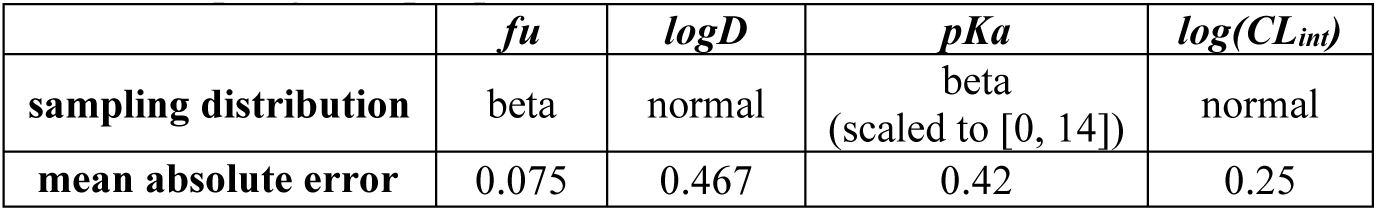
Distribution types and mean absolute errors used for Monte Carlo sampling of input parameters.

#### Model calibration limits *Vss* prediction variance

We began our analysis by examining the variance of *Vss* predictions, as this PK outcome provides direct continuity with the previous fidelity analysis. As there are no experimental *Vss* values to provide ground-truth for our pseudomolecules, the following comparisons are strictly between models. We first examined the variances of the *Vss* distributions produced by propagating identical parameter uncertainty through the four PBPK models. Over the 10^4^ pseudomolecules, median prediction variances were broadly comparable (σ^2^ ∼ 10^-1^, **Table S4**). The one clear exception was the uncalibrated Pearce model, which exhibited a median variance one order of magnitude greater than that of the other three models. Once calibrated, the median variance for the Pearce model fell more in line with the other PBPK models (95% interval 0.16-0.17 L^2^/kg^2^), and this model’s prediction variance became fairly uniform across the pseudomolecule population. Indeed, it appears that model calibration markedly limits the spread of Pearce *Vss* predictions (**Figure S2A**). By contrast, the Mathew model yielded predictions with variance as large as 10^16^ L^2^/kg^2^, depending on the pseudomolecule in question. Notably, there are several outliers with variance greater than 10^7^ L^2^/kg^2^ (**Figure S2B**). The only other model yielding several pseudomolecule outliers was the calibrated form of the Pearce model, which yielded very narrow distributions (∼10^-3^ L^2^/kg^2^) for its outliers – indicative of predictions that are relatively insensitive to parameter uncertainty.

#### Model predictions diverge for a subset of chemical space representing highly protonated, lipophilic molecules

We next quantified the extent to which model predictions agreed under uncertainty and, thereby, sought to identify which regions of chemical space provoke model disagreement. We also broadened our analysis to include dynamic PK outcomes relevant to local tissue exposure. Specifically, we examined *Cmax,u*, *AUCu_0-∞_*, and *T_0.1μM_* in muscle intracellular water. For each pseudomolecule, PK prediction distributions were log-transformed, standardized with a whitening transformation, and contrasted by calculating a Wasserstein distance – a metric that captures differences in both location and shape of paired distributions^86–88^ (see pseudocode in **Algorithm 2**). This analysis produced six distances per PK outcome. Across the four PK outcomes, this resulted in 24 features that summarized model-to-model agreement (predictions from four models, compared pairwise). To distinguish patterns of model agreement across chemical space, we applied K-means clustering to group molecules by these 24 metrics – using a silhouette statistic to select the number of clusters. We thereby identified two groups among the 10^4^ pseudomolecules: a smaller subset (751 molecules; 7.5%) characterized by greater prediction discordance (*i.e.*, larger Wasserstein distances) and a larger subset (9249 molecules; 92.5%) exhibiting broad concordance across models. A heatmap of pairwise distances (**Figure S4**) illustrates the segregation across clusters, and Figure 4A depicts predicted distributions for one prototypical pseudomolecule from each cluster.

**Figure 4.**
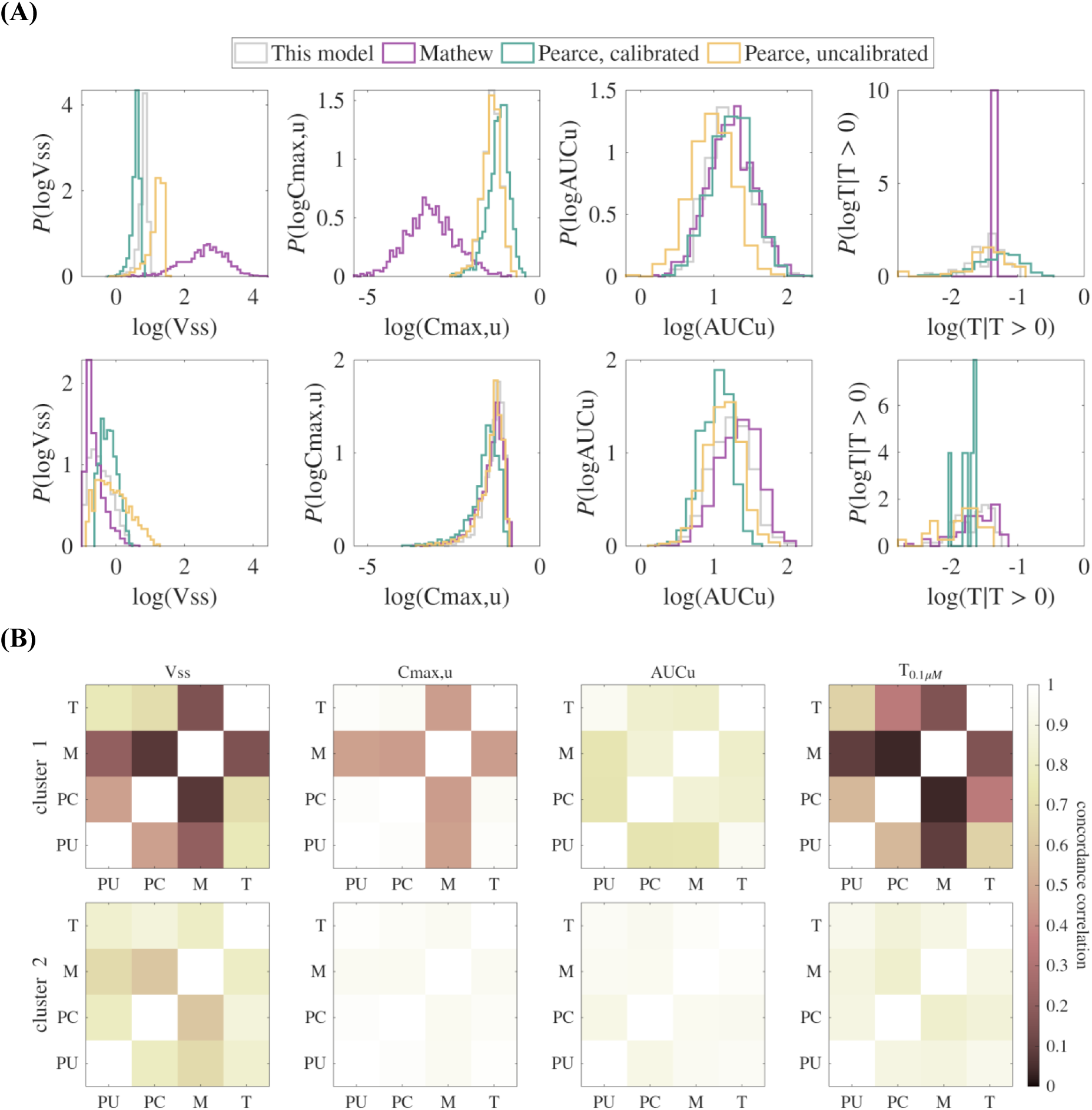
Mathew model exhibits PK outcome prediction discordance for a subset of 751 pseudomolecules (of 10,000 total). **(A)** Distributions of PK outcomes resulting from parameter uncertainty, as predicted by each of the PBPK models. Distributions shown are prototypical of molecules assigned to: Top – Cluster 1. Bottom – Cluster 2. **(B)** Concordance correlation coefficients for predicted PK outcomes, computed across all pseudomolecules in each cluster (Top – Cluster 1: 751 pseudomolecules. Bottom – Cluster 2: 9249 pseudomolecules). Analysis reflects 10^3^ Monte Carlo realizations for each of the 10^4^ pseudomolecules and thus summarizes the outcomes of 10^7^ simulations per PBPK model. Each heatmap shows pairwise agreement between models (T: this model, M: Mathew, PC: Pearce, calibrated, PU: Pearce, uncalibrated).

To quantify the degree of agreement across all molecules within a given cluster, we evaluated the cluster-specific CCC for each PK outcome (Figure 4B). As for the examples shown in Figure 4A **(bottom)**, predicted distributions for molecules in Cluster 2 largely concur across all outcomes (median CCC_cluster 2_ = 0.95; 95%CI:[0.61, 1.00]). The discordance that arises in Cluster 1 (median CCC_cluster 1_ = 0.76; 95%CI:[0.029, 1.00]), is most evident for prediction of *Vss* (CCC_cluster 1,Vss_ = 0.51; 95%CI:[0.083, 1.00]), and *T_0.1μM_* (CCC_cluster 1,T0.1mM_ = 0.40; 95%CI:[0.024, 1.00]). The Mathew model appears to be the principal outlier here, presenting pronounced disagreement with all other models for *Vss*, *Cmax,u*, and *T_0_._1μM_*.

As a first step to determine what distinguishes molecules whose predictions are discordant across models, we examined the biophysicochemical properties of the pseudomolecules assigned to each cluster (**Figure S5**). We found that molecules in Cluster 1 are exclusively bases (78%) and zwitterions (22%), while molecules in Cluster 2 were more evenly distributed across ion classes. Furthermore, as judged by two-sample Kolmogorov-Smirnov tests, molecules in Cluster 1 are more lipophilic (*logD*: D_KS_ = 0.67, Cohen’s d = 1.67) and more likely to be highly protonated (acceptor *pKa*: D_KS_ = 0.70, Cohen’s d = 1.92) than those in Cluster 2 (D_critical_ = 0.063, α= 0.0083; **Figure S6**).

#### For highly protonated, lipophilic molecules, prediction discordance arises from model structure and is exacerbated by parameter uncertainty

We next sought to determine: (1) whether Cluster 1 represents a distinct sub-region of chemical space within which the four PBPK models will make discordant predictions; (2) whether the four models make distinct nominal predictions for molecules in this region or, alternatively, (3) whether the disagreement between models arises due to the simulated parameter uncertainty. Further, if nominal predictions happened to diverge, we established whether one model provided objectively more credible predictions than another. To assess credibility, we needed to compare model predictions to ground-truth, which motivated a return to simulating PK outcomes for molecules in the Obach data set, for which we have empirical data for *Vss*. We performed Monte Carlo simulations for the Obach molecules and used the resulting Wasserstein distance features to map these real drug molecules to the clusters we defined using the pseudomolecules. By examining fidelity of nominal *Vss* predictions alongside the model agreement features, we were able to distinguish whether divergent predictions stemmed from differences in structural assumptions or from the impact of parameter uncertainty.

Twenty-eight percent (N = 240) of the 862 Obach compounds mapped to pseudomolecule Cluster 1 – the cluster corresponding to discordant PBPK predictions under uncertainty. To explore the nature of this discordance, we compared nominal *Vss* predictions between pairs of models. Figure 5 shows a representative result for the two R&R-based models, our model and that of Mathew. Markers indicate the location of Obach molecules, positioned according to the fidelity of their nominal predictions. As the axes report *Vss* prediction fidelity with respect to ground truth, molecules located along the diagonal will be those for which the two models make the same nominal prediction – irrespective of whether that prediction matches the experimentally observed value of *Vss*. The underlying colormap indicates the Wasserstein distance between Monte Carlo-derived *Vss* distributions for the two models.

**Figure 5.**
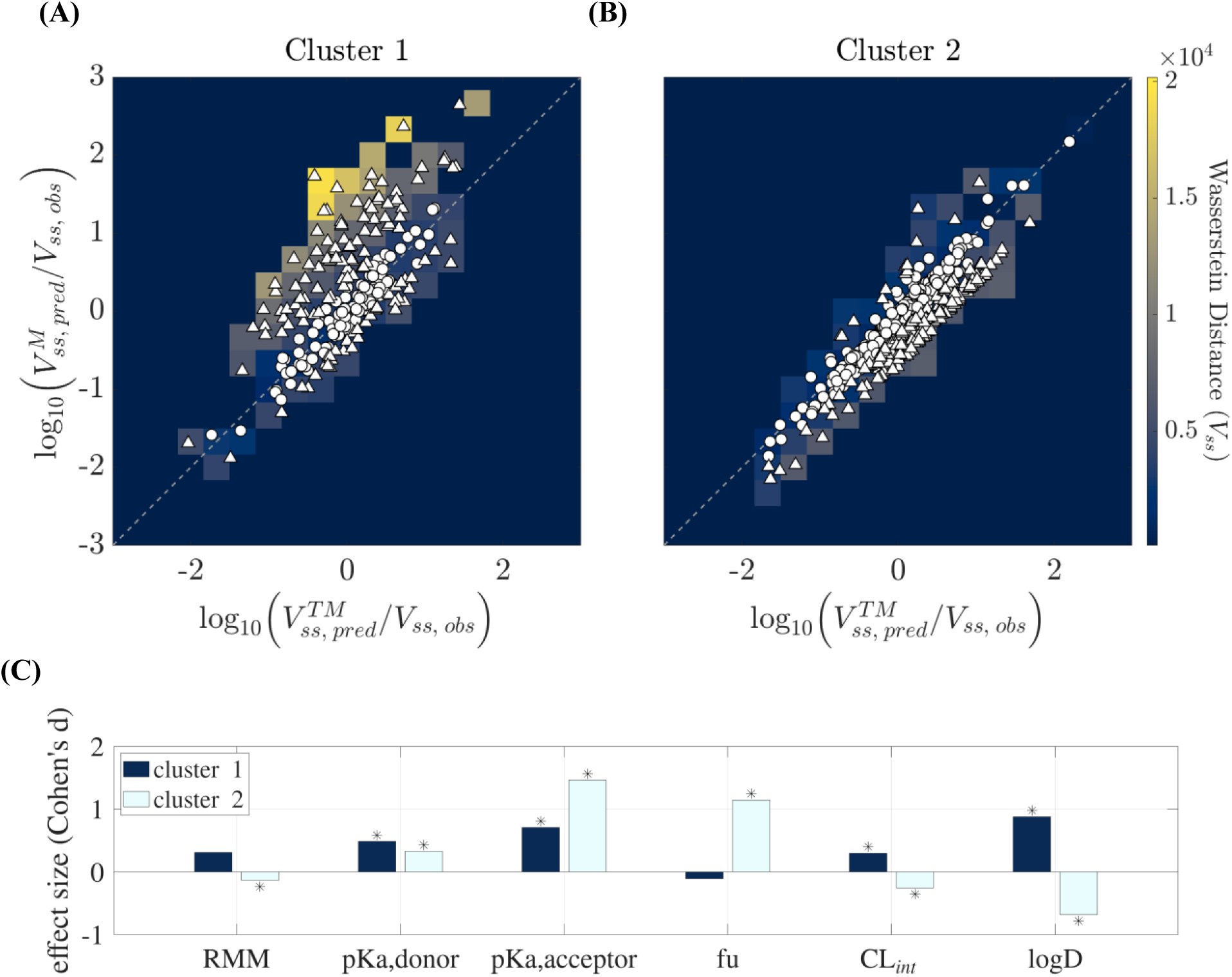
For protonated, lipophilic molecules, the two R&R models yield nominal predictions with greater discordance, and that discordance is exacerbated under uncertainty. Axes report nominal *Vss* prediction fidelity for: X-axis – This model (TM). Y-axis – Mathew model (M). Colormaps indicate the Wasserstein distances between *Vss* distributions predicted under parameter uncertainty. Colormaps are overlaid with markers indicating the locations of individual compounds. Circles: molecules for which the two models’ nominal predictions differ by less than a factor of two. Triangles: compounds for which predictions differ by a factor greater than two. **(A)** Cluster 1: 240 molecules, ≤ 2-fold difference: 108 molecules, > 2-fold difference: 132 molecules. **(B)** Cluster 2: 622 molecules, ≤ 2-fold difference: 397 molecules, > 2-fold difference: 225 molecules. **(C)** Summary of the Cohen’s d effect sizes distinguishing the properties of molecules in each sub-group. Asterisks indicate those features with a statistically significant difference between the near-diagonal and off-diagonal molecules – as determined by Kolmogorov-Smirnov statistic, DKS, exceeding the critical values of Dcritical = 0.21 and 0.14, respectively, for clusters 1 and 2. **Table S6** reports full results of the statistical comparison.

We divided the 240 compounds in Cluster 1 into two groups based on their nominal *Vss* predictions: (a) a near-diagonal set (N = 108), representing compounds whose predictions from the two models differed by less than a factor of two, and (b) an off-diagonal set (N = 132), representing compounds with larger fold-differences in nominal predictions. In Figure 5A, we found – perhaps unsurprisingly – that agreement between nominal predictions (*i.e.*, near-diagonal compounds) appeared to coincide with regions of less discordance under uncertainty – as determined by comparing Wasserstein distances for near diagonal and off-diagonal molecules (Cohen’s d effect size = –1.6, D_KS_ = 0.77, D_critical_ = 0.18). Where nominal predictions disagreed (*i.e.*, off-diagonal markers), the underlying colormap indicated greater discordance under uncertainty. This suggests that much of the observed discordance is due to differences in model structure. Parameter uncertainty did, however, promote greater discordance between models for molecules whose nominal predictions generally concurred (Cohen’s d = 0.94, D_KS_ = 0.35, D_critical_ = 0.15, when comparing Wasserstein distances for near-diagonal molecules in the two clusters). Thus, parameter uncertainty appears to promote greater divergence in model predictions for the type of molecule that appears in Cluster 1.

We next examined the molecular properties of the two subsets within Cluster 1 to assess whether they represent distinct regions of chemical space. Each molecule in the dataset is described by six properties (*RMM, CL_int_, fu, logD,* donor and acceptor *pKa*). Pairwise comparisons of the property distributions for the two subsets, using two-sample Kolmogorov-Smirnov tests, showed statistically significant segregation of medium effect or greater for two properties (D_KS_ > D_critical_ for Bonferroni-corrected confidence level α = 0.0083 and Cohen’s d > 0.5; **Table S6; Figure S7**). In order of effect size magnitude, these were *logD* and acceptor *pKa* (Figure 5C). Accordingly, off-diagonal molecules were more likely to be lipophilic and protonated. A similar analysis of Cluster 2 identified three distinguishing features of medium effect or greater. Off-diagonal molecules in Cluster 2 were characterized by greater acceptor *pKa* and *fu,* but lower *logD*.

Adding these observations to the fact that acceptor *pKa* globally distinguishes clusters 1 and 2 (Cohen’s d = 1.2, α = 0.0083; Figure 6), we posit that the properties typical of molecules whose *Vss* predictions are highly discordant between our model and that of Mathew are: (1) high degree of protonation coupled with (2) high lipophilicity or limited plasma protein binding. Despite their similar underlying framework, the specific assumptions these two models make about the roles of lipophilicity and ionization in determining tissue partitioning produce markedly divergent predictions for molecules of this type. That divergence is thus associated with uncertainty regarding model structure. However, the greater Wasserstein distances observed for the subgroup of molecules for which our model and Mathew’s make more similar predictions indicates that the entirety of Cluster 1 exhibits greater susceptibility to parameter uncertainty than predictions made for molecules in Cluster 2.

**Figure 6.**
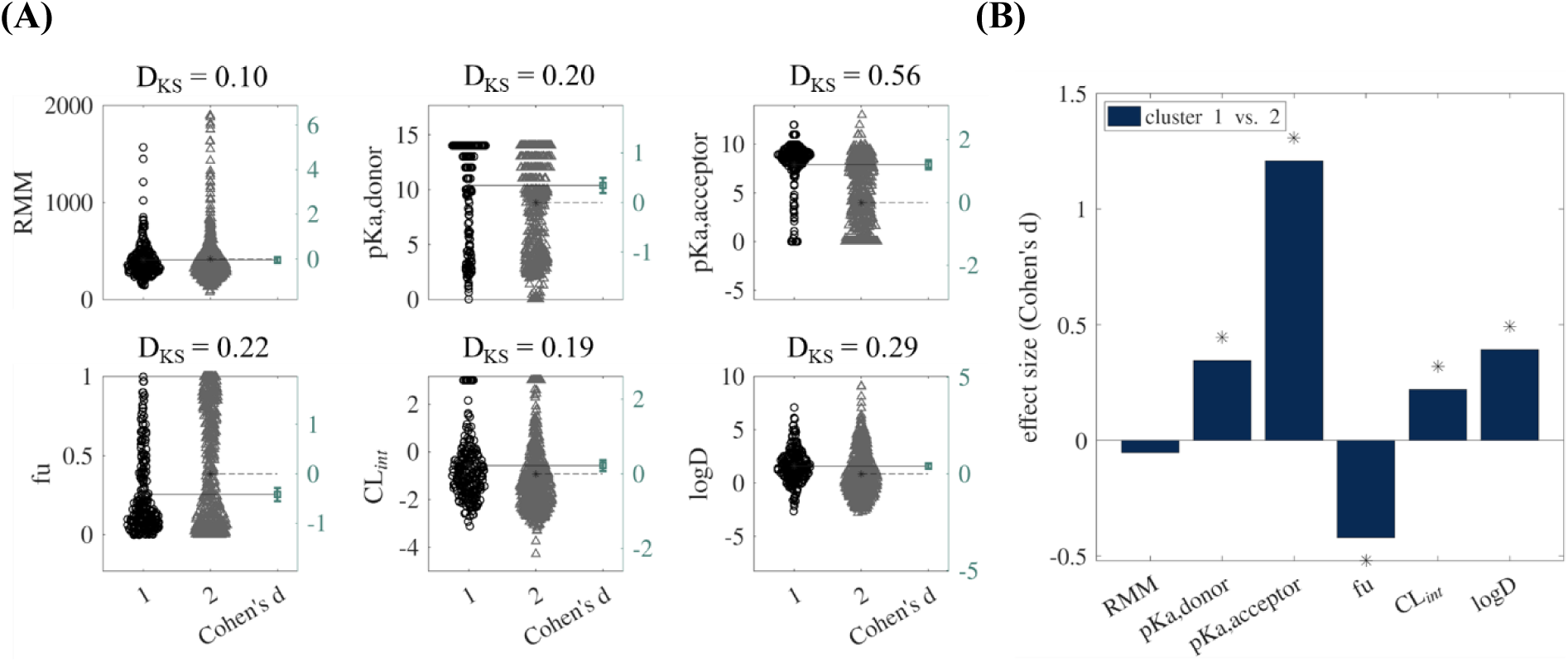
Effect sizes characterizing property differences between Obach molecules assigned to clusters 1 and 2. (A) Gardner-Altman plots of properties for molecules in each cluster. Second y-axis indicates the effect size, calculated as Cohen’s d values, with bootstrapped confidence intervals. (B) Summary of the Cohen’s d effect sizes across all six features. Asterisks indicate those features with a statistically significant difference between the two clusters (as determined by Kolmogorov-Smirnov statistic, DKS, exceeding the critical value Dcritical = 0.13). Number of molecules per cluster: N1=240, N2=622.

#### Subtle changes in model formulation and model calibration markedly impact PK outcome sensitivity to parameter uncertainty

To explore how drug-specific parameters drive prediction variability in the four different models, we performed a Sobol’ global sensitivity analysis. Sobol’ analysis is a variance-based approach that returns indices apportioning the variance of a model’s output between the model’s input parameters. In these analyses, it is common to report the first order and total effect sensitivity indices. These indicate, respectively, the independent parameter contributions to output variance and the total contributions that result from adding the effect of parameter interactions. When input parameters exhibit correlations – as our biophysicochemical properties do (see **Equation 25** for complete correlation matrix) – the interpretation of the sensitivity indices changes, as the correlations between parameter values act as another axis of parameter interaction^89–91^. In this case, the sensitivity analysis can yield indices separating the correlative interactions from the independent interactions.

Using the Monte Carlo estimator for correlated inputs developed by Kucherenko^89^, we evaluated the total correlated sensitivity index, 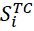, and the total uncorrelated sensitivity index, 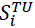, for each of the PBPK model’s drug input parameters. 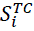 reflects the sum of parameter *i*’s main effect on output variance (*i.e.*, independent of parameter interactions) and the interactions associated with parameter correlations. 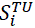 reflects the sum of a given parameter’s main effect and the correlation-independent parameter interactions. Given these definitions, 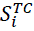 and 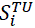 may be greater than or equal to zero, with 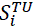 tending to zero as |ρ_*i*,∼*i*_| → 1 (where, ∼ *i* denotes any parameter other than parameter *i*). In the absence of parameter correlations, *i.e.*, |ρ_*i*,∼*i*_| → 0, 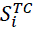 is reduced to the first order sensitivity index, and 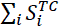 cannot exceed 1. A corollary conclusion is that sums exceeding 1 must reflect correlation effects. Details on our implementation are provided in the Methods section and in the code accompanying this manuscript.

We assessed the properties of drugs in the Obach dataset^68^ and found statistically significant linear correlation coefficients among the six PBPK input parameters. Amongst these, the strongest correlation was between *logD* and *fu* (ρ = −0.56, p < 0.001), followed closely by *logD* and donor *pKa* (ρ = 0.49, p < 0.001). Figure 7 reports the results of the sensitivity analysis for each of the PBPK models and for each cluster identified in the agreement analysis.

**Figure 7.**
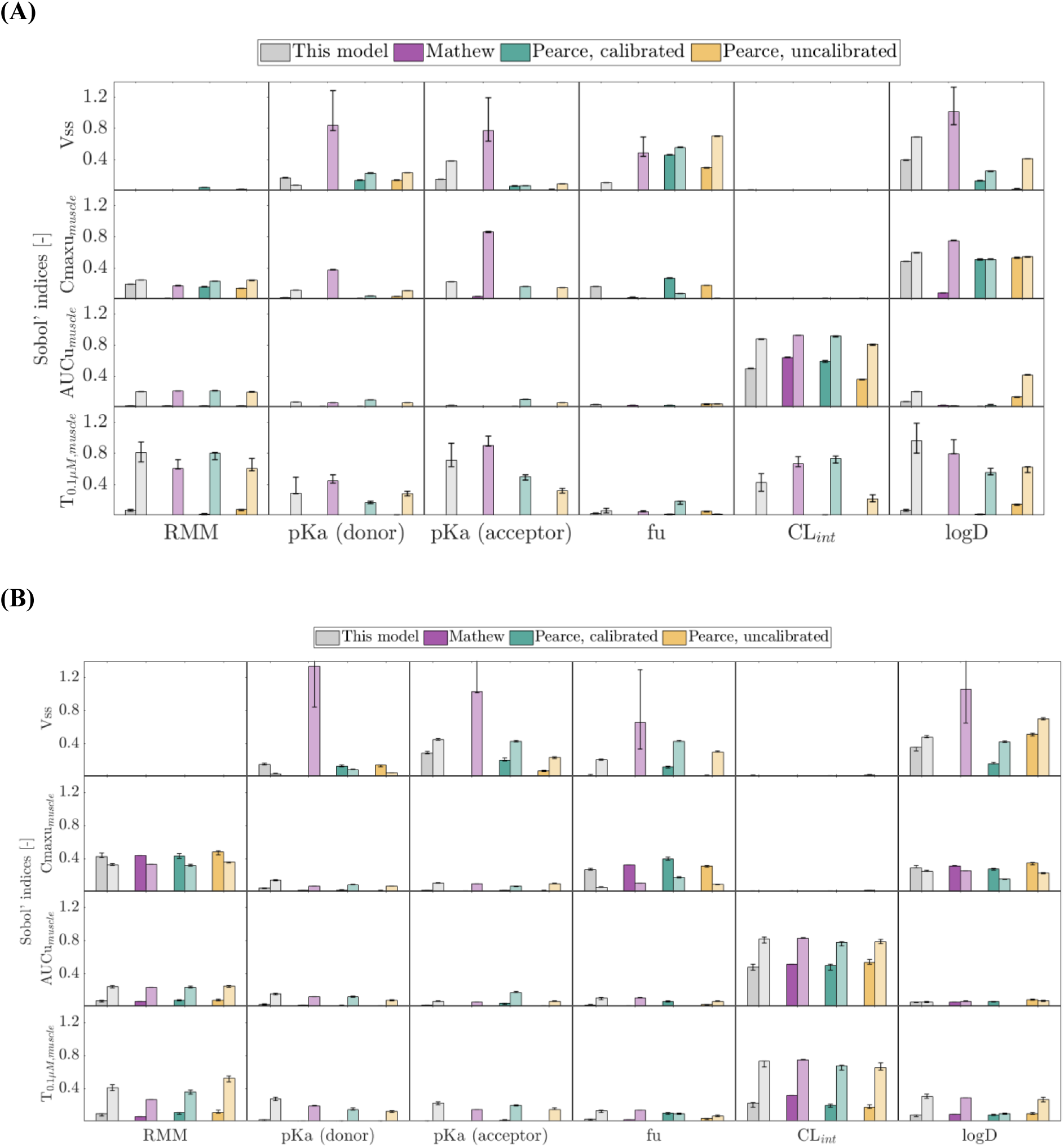
Sobol’ global sensitivity indices for prediction of human PBPK statistics. Paired bars indicate the total correlated (first order + correlative interactions, left) and total uncorrelated (first order + non-correlative interactions, right) indices, respectively. Results are shown separately for property distributions consistent with: **(A)** Cluster 1. Sample sizes for each model were: Nsamples = 1,310,720 (calibrated Pearce); Nsamples = 3,407,870 (uncalibrated Pearce); 11,796,500 (our model); and 12,058,600 (Mathew). **(B)** Cluster 2. Sample sizes for each model were Nsamples = 65,536 (our model and Pearce models); and 25,690,100 (Mathew), respectively. For cluster 2, the upper limits on the confidence intervals for SVss,Mathew are 9.2, 8.4, and 5.6 for donor *pKa*, acceptor *pKa*, and *logD*, respectively. The total number of simulations executed for each model’s Sobol’ analysis was 4×Nsamples. Sample sizes were chosen based on a convergence analysis (see Methods and **Figures S10 & S11**), and error bars indicate bootstrapped 95% confidence intervals.

### Impact of model abstractions on parameter attribution for *Vss*

As lipophilicity is a primary descriptor of tissue partitioning across all models, one might expect its influence on *Vss* to be prominent – perhaps even more so for the subset of chemical space captured by pseudomolecules in Cluster 1, which tend to be of greater lipophilicity (**Figure S6**). However, our model is the only one in which *logD* emerges as the primary driver of *Vss* variance in Cluster 1 (Figure 7A). The influence of *logD* on the *Vss* predictions of the Mathew model is matched or exceeded by that of *pKa*, with a non-trivial contribution from *fu*. By contrast, for Cluster 1, the Pearce model *Vss* predictions are most sensitive to *fu*, and the influence of *logD* appears particularly small in the calibrated model. As the Sobol’ index evaluated here characterizes variability only *within* the interrogated chemical space, and it reflects apportionment of variance rather than absolute variance, these results should not be interpreted as indicating little contribution of *logD* to *Vss* predictions. Rather, the indices tell us that – given the shared lipophilic, protonated nature of molecules in Cluster 1 – the primary drivers of *Vss* variation between molecules shift according to the structure of the PBPK model.

In the case of the Mathew model, this shift exhibits a distinctive qualitative trait, in that the influences of *pKa*, *fu*, and *logD* are entirely attributed to uncorrelated effects. All other models exhibit some degree of correlated sensitivity. This is a direct consequence of the different assumptions each model makes in capturing the role of lipophilicity. Our model and Pearce’s represent lipophilicity by the distribution coefficient, *logD*, which describes the pH-dependent partitioning of neutral and ionized forms of each species. Mathew instead uses *logP*, which does not reflect the impact of ionization. In keeping with this choice, the Mathew implementation limits neutral lipid and neutral phospholipid partitioning to the un-ionized fraction of molecules. These distinct abstractions lead to important differences in the treatment of phospholipid association. As per **Equation 4**, and as demonstrated in **Figure S8**, neutral phospholipid association (*log Knpl)* increases linearly with *logD* in our model (*logMA* = *f* (*logD*)) and that of Pearce (*logMA* = *f* (*logP*)). The Mathew model instead leverages the approximation employed by Rodgers and Rowland^63^ (*Knpl* = 0.3*P*+0.7). This yields a relationship between *log Knpl*, and input parameter *logD* that is non-linear – tending to a constant at low values of *logD* and then rising steeply as *logD* approaches values more typical of drug-like molecules. Consequently, as *logD* increases, the Mathew estimates for *log Knpl* grow to be orders of magnitude greater than the *log Knpl* estimates appearing in the other models.

Whether large estimates for *Knpl* inflate *Kpu* depends on how each model describes the interplay between ionization and lipophilicity. In the Mathew model, neutral phospholipid partitioning only contributes to predicted *Kpu* if molecules are not ionized. In Cluster 1 of the synthetic dataset, only 6% (48/751) of the molecules possess a neutral fraction of > 5%. Thus, for Cluster 1 molecules, the influence of *pKa* prevents large estimates for *Knpl* from impacting *Kpu* through neutral phospholipids. Instead, the effect of *Knpl* will primarily appear via acidic phospholipid (AP) association, which the Mathew model defines as a characteristic of any molecule possessing a protonated site (*i.e.*, cationic and zwitterionic fractions), irrespective of net molecular charge. Eighty-five percent (640/751) of protonatable sites within this cluster of molecules are expected to be highly ionized (defined as having at least 95% of the population protonated at pH 7.4). Coupled with the fact that Cluster 1 molecules tend to be more lipophilic than those in Cluster 2 (*logD*_cluster 1_ = 3.57, 95% CI:[1.56, 7.72] vs. *logD*_cluster 2_ = 0.70, 95% CI:[−2.41, 5.06]), one expects the Mathew model to yield markedly greater predictions for AP association than the other models we considered. Our model and that of Pearce adopt Schmitt’s correlation to describe AP association (**Equation 8**). This formulation allows all species to associate with acidic phospholipids – and to do so with association coefficients that are referenced to *logMA*. By using *logD* to predict *logMA*, our model further tempers AP partitioning for ionized species.

These structural choices explain the Sobol’ results for prediction of *Vss*. In the Mathew model, our sensitivity analysis attributes parameter sensitivity almost entirely to uncorrelated effects across both clusters. This implies the absence of first order effects, as these would otherwise show up as part of the correlated sensitivity index. Instead, each parameter’s influence emerges through non-correlative parameter interactions. The strongest contributors were *logD* and *pKa* – the same parameters whose collective impact we identified as jointly determining AP association. While the other models were also sensitive to these input parameters, the emergent variance observed for *Vss* predictions was split between total correlated and total uncorrelated indices. This pattern recurs for Cluster 2 (Figure 7B). One practical implication of the Mathew model’s strong parameter interactions is that the variance in the predicted PK outcomes cannot be reduced by making high-precision predictions for just the most influential of the drug-specific input parameters. Uncertainty would need to be reduced for all interacting parameters – thereby increasing the performance demands for data-driven property-predictions.

### Model calibration reduces the influence of *logD* and increases that of *fu* on dynamic PK statistics, with notable impact on the relevance of hepatic metabolism

Across all models and both clusters, *Cmax,u* showed negligible sensitivity to intrinsic clearance (*CL_int_*). This is to be expected for a short IV infusion, where the early distribution dynamics are not impacted by hepatic clearance, as would be the case for an oral route of administration. The influence of *CL_int_* instead dominates the variance of *AUCu* and is a pronounced contributor to that of *T_0.1μM_* – both PK statistics that reflect timescales long enough to be influenced by hepatic metabolism.

While *Cmax,u* predictions for molecules in both clusters are largely insensitive to *pKa*, Mathew model predictions for molecules in Cluster 1 again exhibit a pronounced, uncorrelated sensitivity to the ionization parameters and to *logD*. For all other models, *logD* dominates the prediction variance, and the similar magnitudes of correlated and uncorrelated *S_logD_* indices suggest a primarily first-order influence of lipophilicity. For Cluster 2, the Mathew model’s *S_i,cmaxu_* indices mimic those of other models, with correlated indices exceeding uncorrelated sensitivity and variance influences of similar magnitude across *RMM*, *fu* and *logD*.

A difference between the R&R and Pearce model formulations becomes evident in the influences of *logD* and *pKa* on the prediction of *T_0.1μM_* for Cluster 1. Both parameters have a stronger influence on the predictions from the R&R models (This Model and Mathew). Furthermore, in the absence of calibrations, the Pearce predictions of *T_0.1μM_* are relatively insensitive to *CL_int_*, while the R&R models exhibit a pronounced uncorrelated sensitivity. By contrast, predictions of *T_0.1μM_* for molecules in Cluster 2 vary primarily with *CL_int_*, with indices of similar magnitude across all four models. Influences of *logD*, *pKa*, and *fu* are uniformly small.

In addition to inducing *CL_int_* sensitivity for *AUCu* and *T_0.1μM_* predictions in Cluster 1, Pearce model calibrations appear to induce sensitivity to *fu* and reduce the influence of *logD* (Figure 8). This suggests that the core Pearce model tends to overestimate the impact of lipophilicity on tissue partitioning. That effect is then corrected by calibration of *Kp* prior to simulation. Indeed, the Obach compounds whose *Kpu* predictions are most impacted are typically highly lipophilic (**Table S7**), and for these molecules, the tissue most impacted by calibration is adipose (**Table S8**). While reducing *Kpu_adipose_* by ∼93%, model calibrations increased *Kpu_liver_* by ∼42%. Thus, for molecules with high lipophilicity, calibration causes the simulation to render otherwise-inaccessible compounds available for metabolism by reducing hold-up in non-metabolic tissues and increasing exposure in hepatic tissue – producing PK dynamics that respond to *CL_int_* variation. For lipophilic molecules, we thus anticipate that model calibration may create a pronounced qualitative change in PK predictions by ‘gate-keeping’ the relevance of hepatic metabolism.

**Figure 8.**
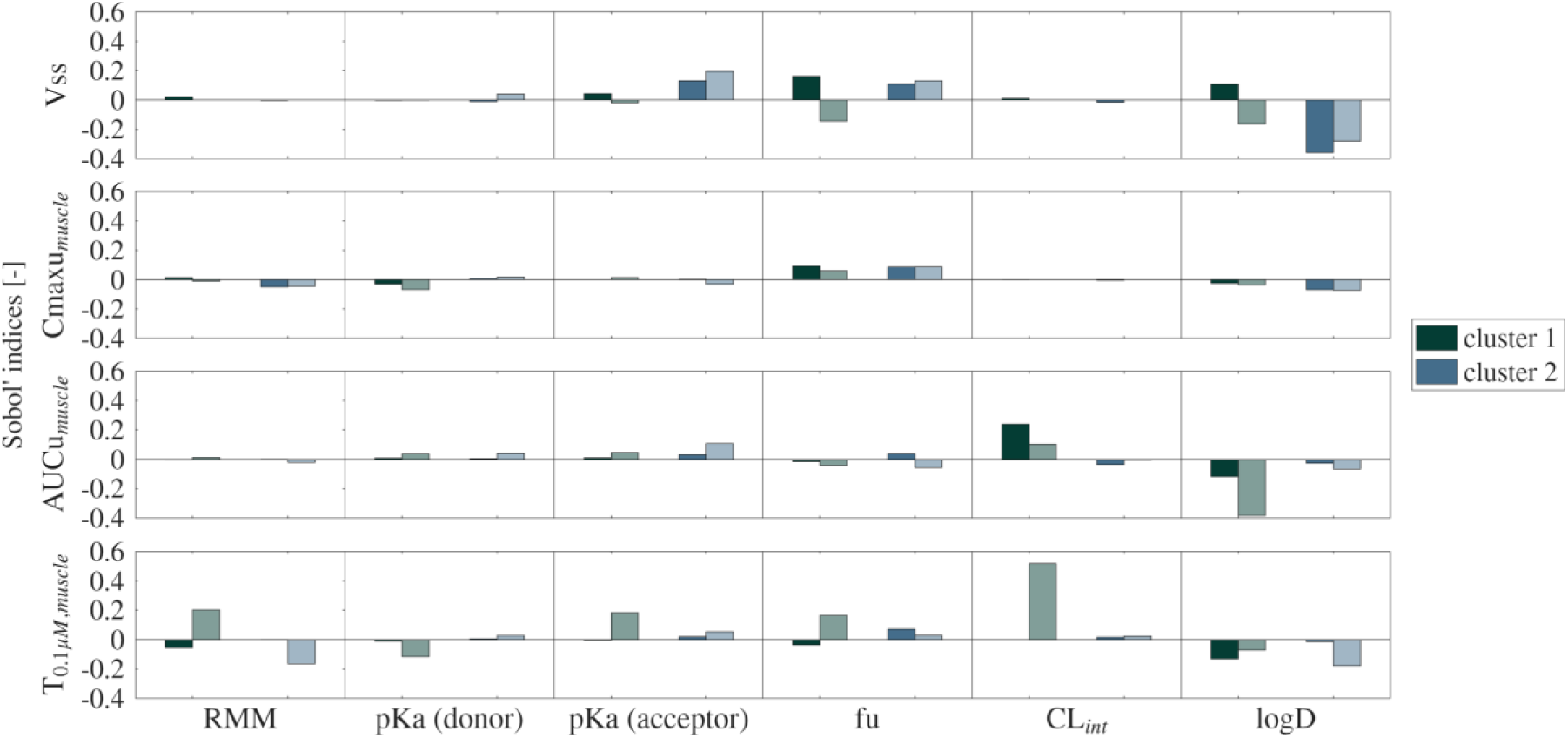
Calibration of the Pearce model reduces the influence of *logD* and increases that of *fu* on the variability of most PK outcomes. Bars indicate how Sobol sensitivity indices change upon calibration of the Pearce model. Paired bars indicate the change in total correlated (first order + correlative interactions, left) and total uncorrelated (first order + non-correlative interactions, right) indices, respectively.

## Conclusion

This study sheds light on challenges to consider in early drug design workflows where input parameters are ML-derived, model assumptions may not be validated, uncertainty is considerable, and decisions may be made without experimental validation. Across experimental datasets, all four PBPK models considered achieved broadly similar fidelity to observed *Vss* and *Kp*, when evaluated with measured input parameters. However, once uncertainty was introduced for the pseudomolecules, clustering of simulation outputs revealed pronounced differences in prediction variance across models and across chemical space. We found that modeling choices that improved nominal fidelity may render models inflexible, thereby risking poor extrapolation to novel regions of chemical space. Modeling choices that invoke ionization state as a filter for lipid and protein association may produce parameter interactions that degrade prediction fidelity and robustness to uncertainty. Furthermore, models tend to diverge in their predictions for protonated, lipophilic molecules, which raises questions about whether any single model will be most credible for decision-making in broad, virtual screens. Our findings do, however, indicate PBPK modeling strategies that may mitigate (or at least avoid exacerbating) issues arising from parameter uncertainty.

### To limit prediction variability, model abstractions should not binarize the inclusion of specific terms as a function of molecule properties

The two R&R models (our model and that of Mathew) illustrate how distinct assumptions within the same nominal modeling framework can lead to very different outputs. The Mathew model consistently exhibited a bias toward underestimation of *Kp* and *V_ss_* and produced greater prediction errors for zwitterions and bases. Classifying a molecule as cationic in the Mathew model heavily weights acidic phospholipid interactions. Furthermore, by adjusting for lipid association in plasma, the assumptions in the Mathew model effectively exclude plasma and interstitial protein binding for protonated molecules that are also classified as lipophilic and highly bound. Consequently, the Mathew model’s predictions are heavily influenced by parameter interactions between *logD* and *pKa*. These modeling choices cause large parameter-driven variance in *Vss* and *T_0.1μM_* once uncertainty is propagated.

### Model calibrations limit prediction flexibility within drug-like chemical space

With respect to *Kp*, the fidelity of the Pearce model is bolstered by calibrating its outputs to a library of pharmaceutical compounds with measured tissue partitioning. However, this calibration has been reported to reduce prediction flexibility as chemical properties vary – a result we also observe herein. Others have postulated this to be an artefact of model’s pharmaceutical-based calibrations being applied to predict outcomes for non-pharmaceutical compounds^92^. Our work suggests that the constraining effect of calibration also limits variability within drug-like chemical space. Alternatively, the calibrations, being based on rat tissue data, may not be appropriate to reflect chemistry-dependent outcome variability in humans. By comparison, our modified model performed comparably to published models, but without the amplified variance observed for the Mathew model or the constrained flexibility associated with the calibrated Pearce model.

### Choice of PBPK mathematical abstraction may frustrate attempts to reduce prediction uncertainty by improving QSAR parameter predictions

Our sensitivity analysis clarified how structural assumptions shape the emergence of variance and pointed to inputs whose refinements would most improve prediction robustness. Variability in *Vss* was driven primarily by *logD* and *pKa*, though their influence was expressed differently across models. For Mathew, variance arose almost exclusively from parameter interactions – indicating that prediction confidence cannot be improved by reducing uncertainty in any single input parameter. Clearer avenues for improvement appear for our model and that of Pearce, as reducing error in inputs such as *fu* or *logD* would directly reduce variance for all PK outcomes. This analysis highlights priorities for improving property prediction models if ML-informed PBPK workflows are to provide informative assessments in virtual screening.

### Limitations

Our analyses included relatively large molecules, which, along with those that are not hepatically metabolized, would violate key model assumptions. Molecular mass did not, however, consistently map to poor *Vss* predictions. In the Obach dataset, of the 101 compounds with RMM greater than 600, only 16 had prediction errors greater than 10-fold in our model’s predictions. Of the 62 total molecules that did have greater than 10-fold prediction errors (26 underpredicted and 36 overpredicted), many are chemotherapeutic compounds that intercalate with DNA (effectively, a target-mediated disposition factor), antibiotics and natural products that are not metabolized, or compounds that are metabolized by intracellular processes that are not specific to the liver. Others partition extensively into blood cells in a way that is not captured by the partitioning phenomena addressed in our generic model – such as binding to cytosolic carbonic anhydrase^93^. A previously reported example of this latter issue, identified by Rodgers and Rowland^94^, is chlorthalidone, for which our model underpredicts *Vss* by ∼8-fold. This differs from the substantial overprediction that would result had we used the commonly employed approach of inferring *K_APL_* from blood cell partitioning. In that case, we would have incorrectly attributed the high level of chlorthalidone association with blood cells to the presence of acidic phospholipids. Propagating that error through to *Kpu* predictions for all tissues would lead to >10-fold overestimation for *Vss*.

We must bear in mind that the fidelity evaluation herein employs point estimates for both the empirical data and our model predictions. We note that, for some drugs, there are discrepancies between the *Vss* metrics reported in Obach *et al.*^68^ and findings elsewhere in the literature – cases where the alternative empirical value is within 10-fold of our model prediction.

Finally, the conclusions presented herein are particular to the IV route of administration simulated. Similar analyses would be required to determine model performance under uncertainty for other routes of administration. For example, *CL_int_* is expected to strongly influence *Cmax,u* for an oral dose, due to first-pass elimination and interplay of absorption kinetics with tissue disposition.

## Methods

Table 1 summarizes the distinguishing features of the three *Kp* models. These features and relevant governing equations are elaborated upon in the section that ows.

### Model implementation

#### General equation for R&R-Lukacova models

Equation 1 captures the decomposition of PBPK partition coefficients into contributions from tissue water, interstitial proteins, and lipid components of each tissue.

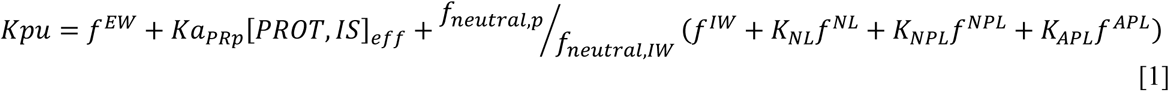

Here, *f*^*i*^ indicates the fractional volume of tissue that is occupied by component *i*, and *f*_*j*_ indicates the micro-ionization fraction of drug in charge state *j*. Plasma is denoted by the subscript *p*, and *Ka_PRp_* denotes the binding affinity for plasma proteins. We denote tissue components as: extracellular water (EW), intracellular water (IW), neutral lipids (NL), neutral phospholipids (NPL), acidic phospholipids (APL), lipoproteins (LPP), albumin (ALB), and intracellular proteins (PROT). Binding to interstitial proteins is expressed as a function of the effective concentrations of those proteins relative to the protein content of blood plasma, [*PROT,IS*]*_eff_*. We denote the association coefficient for each tissue component as *K_i_*.

#### Neutral lipids (plasma and intracellular)

The R&R-Lukacova approach typically employs lipophilicity, quantified by the octanol-water partition coefficient, *logP_ow_*, to predict drug partitioning into neutral lipids. These models capture the consequences of ionization by weighting this term by the fraction of drug molecules expected to remain in the neutral state. In our model, we instead follow the example of Schmitt, and use the pH-specific distribution coefficient, *logD_ow_*, to represent neutral lipid partitioning.

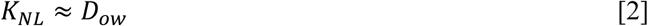

We address the difference in plasma pH and reported differences in intracellular pH across tissues by adjusting *D_ow_* to reflect local states of micro-ionization. Using the Henderson-Hasselbalch equation, we predict the fraction of molecules in neutral and charged states at the relevant local pH. The neutral fraction then informs our distribution coefficient estimates, as indicated in **Equation 3**, where α=0.001 ^95^.

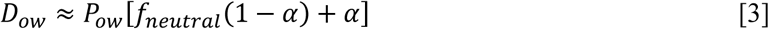

#### Neutral phospholipids

We also apply this local distribution coefficient to predict neutral phospholipid association of ionizable species. Our approach follows that of Pearce *et al.*^60^, who reported improved *Kp* predictions when using the membrane affinity correlation of Yun and Edginton^96^ to approximate neutral phospholipid partitioning (**Equation 4**). Herein, we have assessed the further impact of replacing *logP* with the local distribution coefficient, *logD*, as a potential approach to account for charge effects.

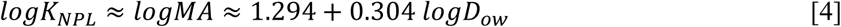

where *MA* is the membrane affinity constant. Finally, we note the adjustment made to account for the triglyceride rich composition of neutral lipids in adipose tissue. As with prior models, we replace the octanol-water partition coefficient, *P_ow_*, with the vegetable oil-water partition coefficient, *P_vow_,* when modeling tissue partitioning for adipose tissue – estimating *P_vow_* from *P_ow_* as per **Equation 5**.^97^ Converting *logP_vow_* to *logD_vow_*, using **Equation 3**, provides the required value for *K_NL_* in adipose tissue.

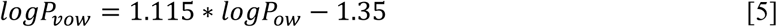

#### Intracellular proteins

The R&R-Lukacova framework does not typically reflect intracellular protein binding^62,63,98^. For our model we add this component to the general equation as:

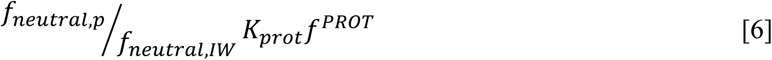

with the first term denoting the charge-dependent difference in concentration between unbound drug in the plasma and intracellular water. Following the example of Schmitt^95^, we approximate the non-specific protein binding coefficient as a function of the phospholipid partitioning coefficient (**Equation 7**).

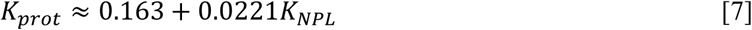

#### Acidic phospholipids (plasma and intracellular)

The R&R-Lukacova framework infers acidic phospholipid partition coefficients from empirically assessed blood-to-plasma ratio and treats the association with acidic phospholipids as a site-specific, 1:1 binding event. Here, we instead adopt the approach of Schmitt and project the affinity for acidic phospholipids as a perturbation of the affinity for neutral phospholipids. Due to electrostatic effects, the cationic fraction of drug is expected to augment the apparent membrane association, and the anionic fraction is expected to reduce that affinity. Based on the correlation proposed by Schmitt^95^, Pearce *et al.*^60^ report the corresponding estimate for *K_APL_* as:

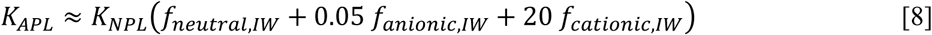

For zwitterions, we adopt the assumption stated by Rodgers and Rowland – namely, that interactions between protonated sites and acidic phospholipids dominate^62^. Thus, we add the zwitterionic fraction, *f*_*Zwitterionic*,*IW*_ to the cationic term in **Equation 8**. Note that this differs from the approach implemented in the high-throughput toxicokinetic model Httk, as part of the framework originally reported by Pearce^60^. There, the zwitterionic fraction was combined with the neutral fraction.

#### Binding to plasma and interstitial proteins

As in the work of Rodgers and Rowland, we use the apparent binding coefficient between plasma proteins and drug to approximate binding to tissue interstitial proteins. If plasma protein binding is empirically quantified via the measurement of unbound drug in plasma samples (*fu*), the apparent binding coefficient can be calculated using **Equation 9**.

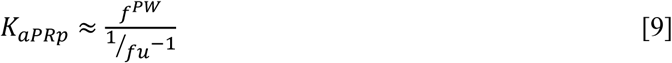

where PW indicates plasma water. This equation assumes that the drug sequestration observed in plasma binding assays is dominated by protein interactions.

As is common practice, we adjust this value to account for the potential impact of drug partitioning into plasma lipid components. Here, we use the adjusted form suggested by Lukacova *et al.*^98^, modified to account separately for neutral lipid, neutral phospholipid, and acidic phospholipid components of plasma.

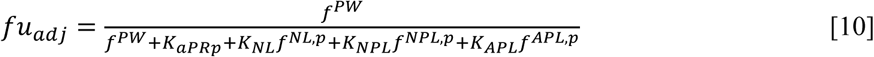

Finally, we capture binding to interstitial albumin and lipoproteins by weighting the apparent ratio of those proteins’ contribution against the estimated binding propensity of albumin and lipoproteins in plasma. As neutral compounds are expected to associate with lipoproteins and anionic components are expected to interact with albumin^99,100^, each protein’s contribution to interstitial binding is weighted by the relevant micro-ionization fraction (**Equation 11**). Here, we have assumed that zwitterions follow the behavior of anions, interacting with albumin.

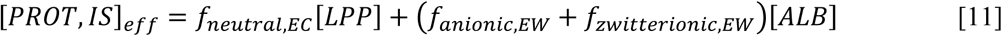

### Prior Models

#### Pearce model

Our implementation of the model reported by Pearce *et al.* closely follows the approach described in their 2017 publication and laid out in the Httk toxicokinetic R package.^60,66^ The general form of their tissue partitioning equation is captured in **Equations 12-15**, and key distinguishing assumptions of the model are detailed in Table 1. The *Kpu* equation has 2 broad components, one term each for interstitial and intracellular partitioning.

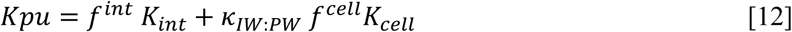

where *int* indicates tissue interstitial space, *f^int^*and *f^cell^* are the fractional volumes of the interstitial and cellular spaces, and *K_int_* and *K_cell_* are the net partition coefficients for each of these compartments (with respect to plasma water). κ_*IW*:*PW*_ captures the impact of the ionization gradient that arises from the difference between intra– and extra-cellular pH, as determined by **Equation 13**.

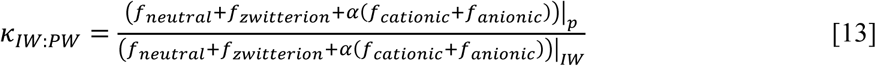

The interstitial partition coefficient, *K_int_*, reflects drug dissolved in interstitial water and non-specific protein binding as follows.

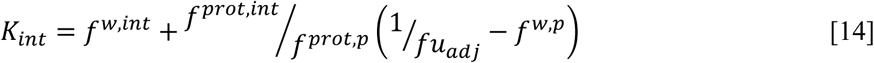

where *w* indicates water, *f^w,int^* and *f^prot,int^* are the fractional volume of water and protein in the interstitial space, and *f^w,p^*and *f^prot,p^* are the corresponding values for blood plasma. The intracellular partition coefficient, *K_cell_*, reflects drug dissolved in intracellular water, along with lipid and protein binding.

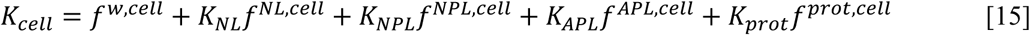

All superscripts with ‘cell’ are fractional volumes expressed relative to cell volume. *K_NL_* and *K_prot_* follow the same assumptions captured in **Equations 2 and 4**, which our model has adopted from that of Pearce. Note that, for all drugs, we used the correlation of Yun and Edginton as a surrogate for predicting membrane affinity, which serves, in turn, as a surrogate for neutral phospholipid partitioning. This differs from the approach reported by Pearce, which employed empirical values for *logMA*, where available.

Finally, this model includes a correction to adjust the *fu* measured *in vitro* to account for the role that lipid partitioning is expected to play in augmenting the apparent bound fraction (**Equation 16**).

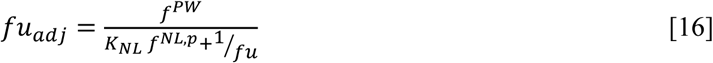

#### Mathew model

Our implementation of the model reported by Mathew *et al.* follows the governing equations made available in the VBA code included with the supplement to that publication^59^. However, we made two key modifications. Firstly, we corrected the equation used to adjust the unbound fraction of drug in plasma, as per Equation 17.

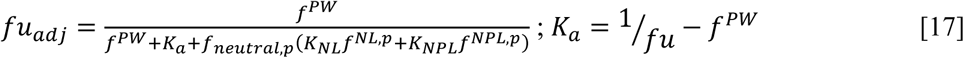

This addresses a sign error that adjusted *fu* upward instead of decreasing its value, as is necessary to account for postulated lipid partitioning. This change obviated the need for the upper limit on *fu_adj_* that the authors imposed. Secondly, we sought to remove the model’s reliance on blood-to-plasma ratio as an input parameter. Our motivation for this is two-fold: (1) the molecule datasets used in this study do not include empirical values for blood-to-plasma concentration ratio (*B2P*); and (2) this parameter proves challenging to predict via QSAR models^101^, and its uncertainty has pronounced first-order impacts on the prediction of human volume of distribution^102^. In lieu of using *B2P* to infer association coefficients for acidic phospholipids, we adopted the Schmitt-Pearce approach and predicted *K_APL_* from the charge-based modification of *K_NPL_.* In Table 1, we further highlight distinguishing assumptions of the Mathew model.

### PBPK simulations

Our 14-compartment PBPK models (**Figure S9**) assume perfusion-limited disposition following a 15 mg/kg IV dose simulated over 1 minute. We simulated disposition dynamics over a time span of 48 hours. For elimination, the models assume hepatic metabolism (via the well-stirred approximation) and passive renal filtration. As such, the models neglect active transport and permeability limitations. They also do not account for target-mediated disposition or non-linear PK phenomena. For example, by using first order kinetics to describe hepatic clearance, the models neglect any saturation of metabolic capacity.

The dynamic PK statistics selected for analysis were the area under the unbound concentration-time curve (*AUCu_0-∞_*), maximum unbound concentration (*Cmax,u*), and time for which unbound concentration exceeds a threshold of 0.1 μM (*T_0.1μM_*). In all cases, the analysis focused on the predicted unbound concentration in the muscle intracellular space.

### Definition of ion classes

When classifying molecules, we defined zwitterions as any compound possessing both protonatable and de-protonatable sites, irrespective of the ionization state at physiological pH. Bases and acids were those with only an acceptor site and those with only a donor site, respectively. Compounds without ionizable sites were classified as neutral. For our synthetic dataset of pseudomolecules, we defined sites to be ionizable if *pKa_donor_* < 9.4 or *pKa_acceptor_* > 5.4 – each of which maps to 1% ionization at pH of 7.4.

### Evaluation of model performance

To assess the prediction fidelity, relative performance, and uncertainty associated with each model, we predicted *Kp* and *Vss* quantities for which empirical data have been reported. *Kp* represents the ratio of tissue concentration to that of blood plasma, and is related to *Kpu* by:

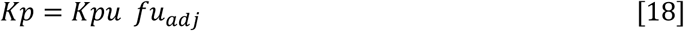

*Vss*, the steady-state volume of distribution, reflects the extent to which a drug is expected to distribute beyond the blood plasma into tissue matrices. Estimation of *Vss* based on mechanistic predictions of *Kpu* is carried out using **Equation 19.**

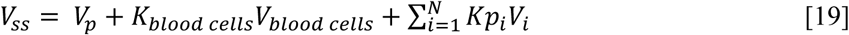

where *V_i_* represents tissue volume, and the sum runs over all tissues.

We used the following statistics to assess model fidelity to data:

- fraction of drugs with PK predictions within 2-fold, 3-fold, and 10-fold error

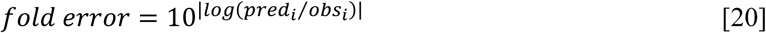
- root mean squared error of log-transformed values

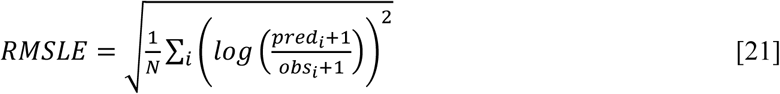
- average fold error

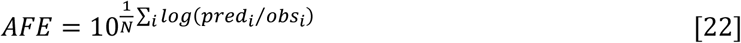
- concordance correlation coefficient (of log-transformed values).

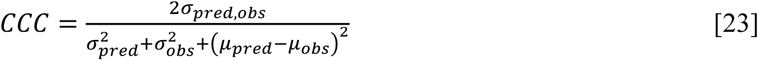

### Monte Carlo simulations

To evaluate the local variance of model predictions subject to parameter uncertainty, we generated pseudomolecules from the Obach dataset by correlated sampling of the key biophysicochemical input parameters *RMM*, *fu*, *logD*, *pKa*, and *CL_int_*. To achieve this, we first assumed a uniform input distribution for each parameter and then imposed correlation by applying a Gaussian copula derived from the property distributions reflected in the Obach dataset (using MATLAB function *copulafit*). We thereby generated a dataset of 10^4^ pseudomolecules, each defined by *RMM*, *fu*, *logD*, *pKa*, and *CL_int_*.

For each pseudomolecule, we then perturbed its properties by sampling from a distribution with mean equal to that of the molecule and mean absolute error matched to the predictive accuracy of typical QSAR models^56–58^. **Table 2** reports the corresponding distribution types and typical errors.

To evaluate the effects of parameter uncertainty, we simulated 1,000 Monte Carlo realizations for each of 10^4^ pseudomolecules via each of the four PBPK models of interest – giving a total of 4×10^7^ simulations. Each realization predicted four PK outcomes: steady-state volume of distribution (*Vss*), maximum unbound concentration (*Cmax,u*), area under the unbound concentration-time curve (*AUCu*), and the duration of time for which concentration exceeded 0.1 µM (*T_0.1µM_*). The latter three were generated for intracellular concentrations in the tissue of interest.

### Pseudocode for generation of synthetic pseudomolecules and parameter perturbations for Monte Carlo simulations

**Algorithm 1.**
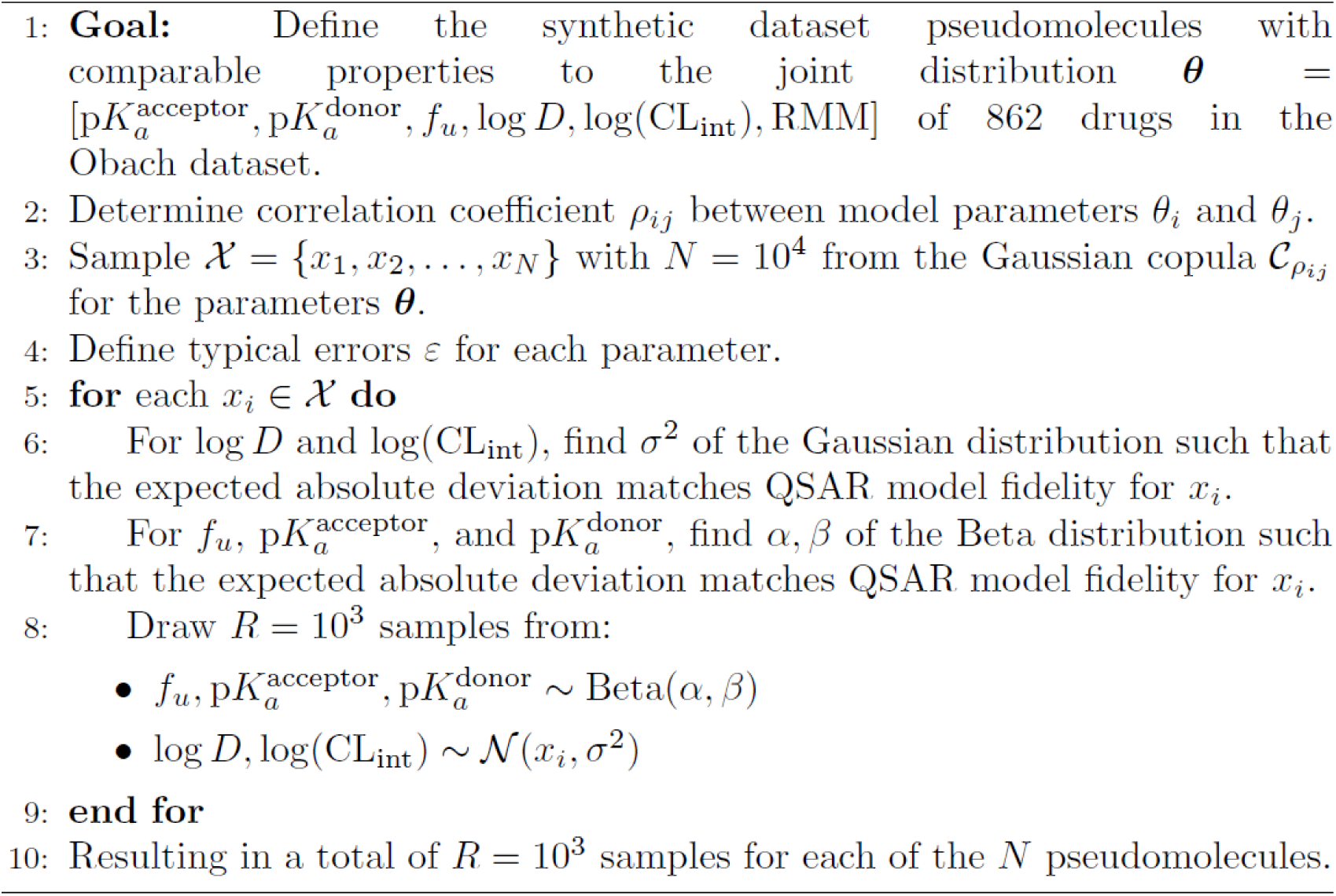
Generation of Synthetic Pseudomolecules.

### Model agreement analysis

The Monte Carlo simulations yielded distributions of the four predicted PK outcomes for each molecule and each PBPK model. We used these distributions to assess the extent to which models generate predictions that deviate under parameter uncertainty. To this end, we quantified the model-to-model Wasserstein distance between the predicted distributions for a given PK outcome. For each molecule (N_pseudomolecules_ = 10^4^), given four models (N_models_ = 4) and four PK outcomes (N_PKoutcomes_ = 4), this yielded 24 Wasserstein distance features (as per **Equation 24** and **Figure S3**).

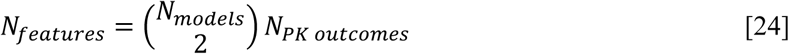

We then used these 24 features to group molecules into consensus clusters based on model agreement patterns. The PK preprocessing applied before distance calculation and the feature transformation applied prior to clustering are described in detail in **Algorithm 2**. Also detailed is our approach to defining robust cluster assignments. We performed 1000 K-means clustering attempts, noting with each attempt which pairs of molecules were assigned to the same group. By tallying co-clustering events, we were able to identify groups of molecules that clustered together in at least 95% of the clustering attempts. This approach removes any molecule that does not meet the 95% frequency threshold for stably clustering with other molecules. For this dataset, however, we found that all molecules could robustly be assigned to a cluster.

In addition to our whitened/PCA-transformed analysis, we evaluated clusters defined by omitting the preprocessing step (*i.e.*, calculating Wasserstein distances based on raw PK data) and/or by applying min-max or principal component feature transformations. The final number of clusters (2 or 3) – and their size – varied with the choice of pre-processing and feature transformation approach (see adjusted Rand indices, **Table S5**). We arrived at the decision to employ whitening and PCA transformation by examining the silhouette statistic distributions resulting from each approach (**Table S5**). We also considered the potential of whitening decorrelation to partially segregate global and local influences on tissue exposure (*i.e.*, bulk changes in exposure to all body tissues that might drive increases in *Vss* vs. local changes in muscle intracellular exposure that are driven by the interplay of muscle perfusion and tissue composition).

### Pseudocode for whitening transformation and K-means clustering

**Algorithm 2.**
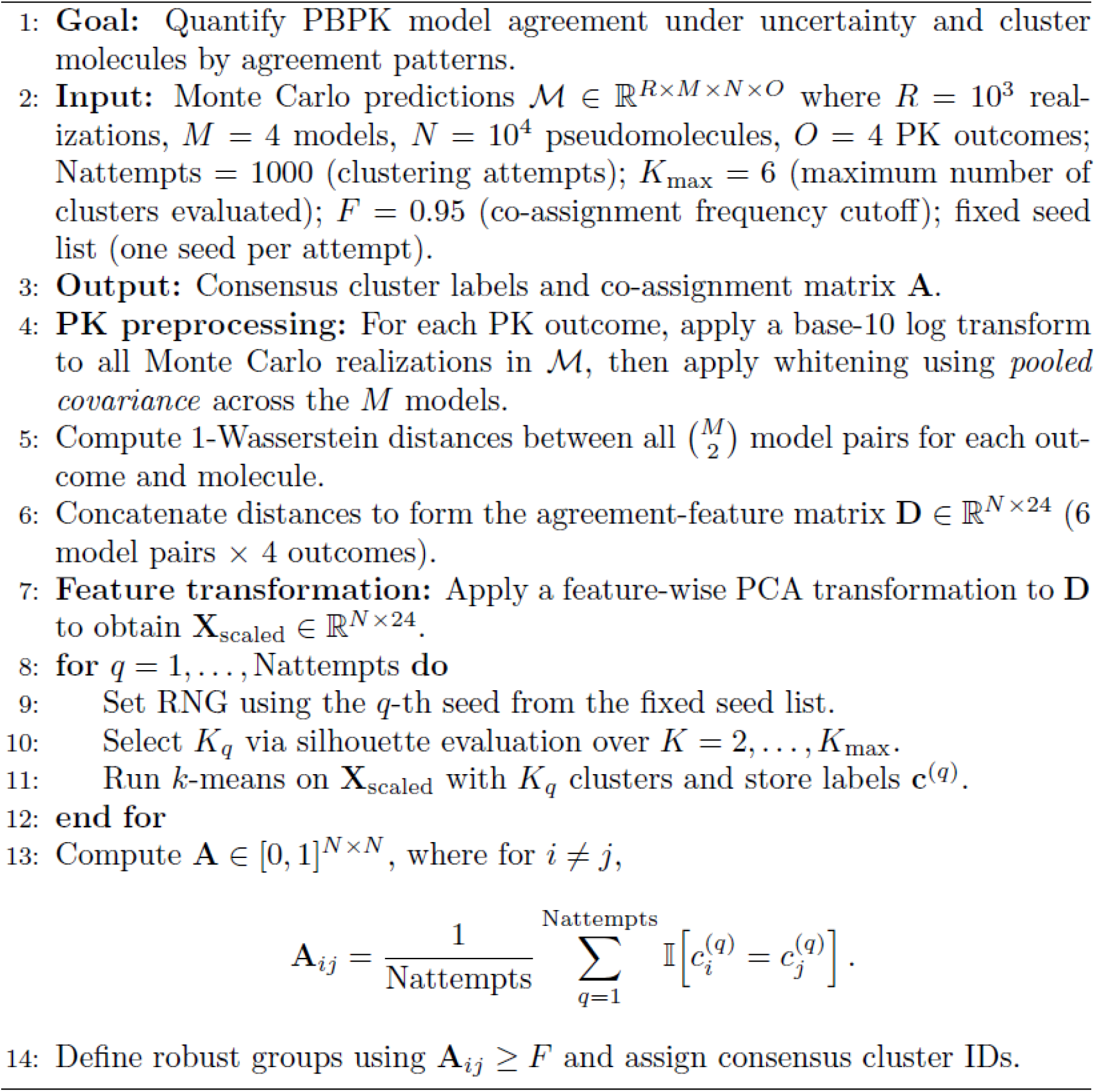
Model agreement analysis: PK preprocessing and feature scaling.

### Obach molecule discordance analysis

To connect the cluster patterns from the pseudomolecule analysis to real drug space and distinguish structure-driven disagreement from the effects of parameter uncertainty, we analyzed 862 molecules from the Obach dataset. We performed Monte Carlo simulations for these molecules and computed the 24-element Wasserstein distance features required to map these real molecules to the feature-space defined by our synthetic dataset. This required whitening and PCA transformations using the same transformation matrices obtained in our pseudomolecule analysis. We then assigned molecules to clusters by identifying which cluster centroid was closest to each Obach molecule’s feature vector (as defined by Euclidean distance).

#### Statistical analysis of sub-group properties

After mapping molecules to clusters, we used two-sample Kolmogorov-Smirnov (KS) tests to assess whether these groups corresponded to distinct sub-regions of chemical space. We applied this analysis to the six biophysicochemical properties of interest across the two clusters, using a Bonferroni correction to adjust the chosen confidence level of α = 0.05 for these multiple comparisons. For all comparisons, we also quantified the Cohen’s d effect size associated with property differences between groups. A similar workflow was used to determine any differences between cluster sub-groups corresponding to ≤ 2-fold and >2-fold differences in *Vss* values predicted by our model and that of Mathew.

### Sobol sensitivity analysis

We performed variance-based global sensitivity analyses using the approach described by Kucherenko *et al.* for correlated input parameters^89^. The statistically significant correlation coefficients for molecules in our chosen subset of 862 molecules from the Obach dataset are presented as **Equation 25**, with the range of drug property values indicated in **Table 3**.

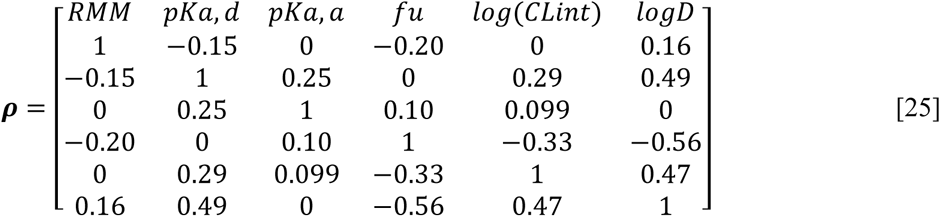

**Table 3.** 95% interval of drug property values for selected molecules in the Obach dataset.

|  | RMM | pKa, donor | pKa, acceptor | fu | log(CLint [L/kg]) | logD |
| --- | --- | --- | --- | --- | --- | --- |
| **lower bound** | 153 | 0.713 | 0 | 0.002 | -2.81 | -2.20 |
| **upper bound** | 1160 | 14 | 11.0 | 1 | 3 | 5.03 |

#### Convergence analysis

For the Sobol’ analysis, we assumed a uniform input distribution for each parameter and imposed correlation by applying a Gaussian copula derived from the property distributions reflected in the Obach dataset (using MATLAB function *copulafit*). We confirmed convergence of the Monte Carlo estimator by assessing the 95% interval of sensitivity indices estimated from 100 bootstrapped samples. As a convergence statistic^103,104^, we monitored the largest 95% estimation interval characterizing predicted PK statistic *j*, observed over all *n* parameters, as per **Equation 26**.

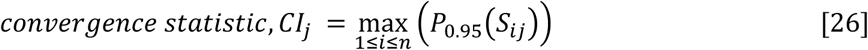

In most cases, we evaluated sufficient samples to reduce this interval to << 0.1 (**Figures S10 & S11**). A notable exception is the case of *Vss* prediction by the Mathew model, where the Monte Carlo estimator had not converged after running ∼2.6×10^7^ samples (requiring >10^8^ simulations).

To validate that apparently influential parameters could be distinguished statistically from those of negligible influence, we implemented a dummy model output to act as a control. This output’s value was set equal to a uniform random input parameter (*i.e.*, a parameter that did not feature in the PBPK model and was not correlated with any other parameter). When defined this way, the dummy output acted as a positive control for true, first order sensitivity to one parameter 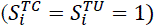 and a negative control reflecting true insensitivity to all other parameters 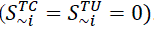. A parameter was then considered to be influential if the median of its bootstrapped index estimates differed from that of the negative control (*i.e.*, medians were determined to be statistically distinct by a Wilcoxon rank-sum test at confidence level of α = 0.05).

### Programming languages and code availability

We performed all simulations and analyses in MATLAB (The MathWorks Inc., Natick, Massachusetts). Differences between distributions were computed using the Wasserstein distance metric, as implemented in the *ws_distance* function^105^, and covariance normalization was performed using ZCA whitening as implemented in the *whiten* function^106^. Differences in clustering outputs across preprocessing and feature-standardization approaches were quantified using an adjusted Rand index, computed using function *randindex*^107^. Seeds for random number generation are detailed in **Table S9**.

Details of our computing environments appear in **Table 4**. Data and code used to perform the analyses for this paper are hosted at https://github.com/MADBioLab/PBPKPredictionVariability.

**Table 4.**
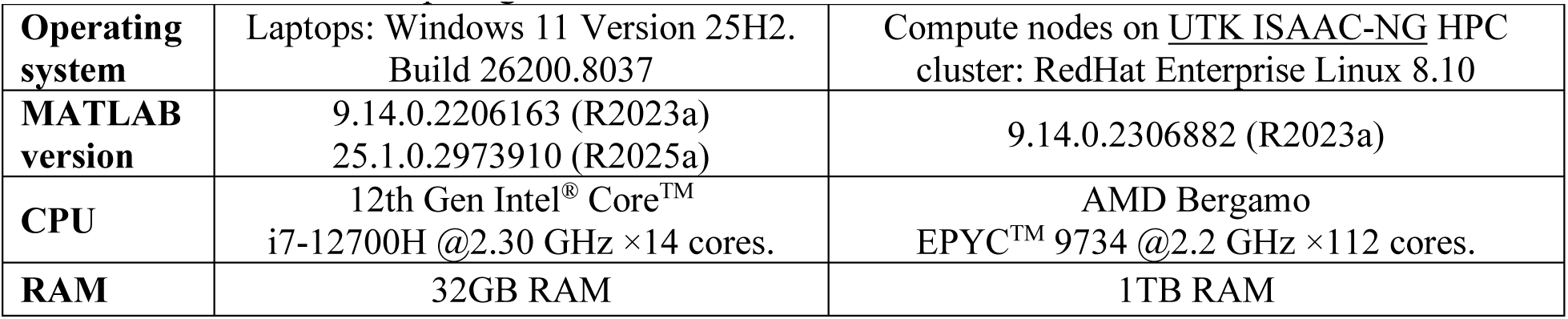
Software and computing environment details.

## Acknowledgements

A portion of the computation for this work was performed on the University of Tennessee Infrastructure for Scientific Applications and Advanced Computing (ISAAC) computational resources. BSA acknowledges the Accelerating Therapeutics for Opportunities in Medicine (ATOM) Consortium. The authors would like to thank MADBio Lab member Ghizelle Abarro for her comments on the manuscript.

## Author Contributions (CRediT Taxonomy)

Investigation: BSA, MF

Conceptualization: BSA

Funding Acquisition: BSA

Writing-original draft: BSA, MF, ZRF

Writing-review and editing: BSA, MF, DTF, ZRF

Methodology: BSA, MF, DTF

Software: BSA, MF, DTF

Formal Analysis: BSA, MF, ZRF

## Materials & Correspondence

Belinda S Akpa

