## Supplemental Tables and Figures for "Prediction variability in physiologically based pharmacokinetic modeling of tissue disposition under deep uncertainty"

### Supplementary Information

**Table S1. Dataset provenance**

| Dataset | Source | Scope (size) | Role in this study |
|---|---|---|---|
| **Physiological parameters** | Pearce *et al.*[1] (httk: physiology.data) | Human and rat parameter table | Physiological inputs for all dynamic PBPK simulations |
| **Tissue vascular fraction** | Edginton *et al.*[2] | Literature constants | |
| **Tissue composition** | Pearce *et al.*[1] (httk: tissuedata.data) | Human and rat parameter tables | Composition inputs for *Kpu* calculations |
| **Tissue albumin and lipoprotein concentration** | Rodgers *et al.*[3] | Literature constants | |
| **Experimental tissue partitioning coefficients, rat (*Kp*)** | Pearce *et al.*[4] | 157 compounds, 11 tissues. N = 992 total datapoints | Validation datasets for fidelity analyses |
| **Human volume of distribution (*Vss*)** | Obach *et al.*[5] | N = 862 compounds (subset with complete PBPK inputs) | |
| **Empirical biophysicochemical parameters** | Obach *et al.*[5] | N = 862 compounds (subset with complete PBPK inputs) | (1) Drug-specific inputs for PBPK modeling of real compounds. (2) Copula construction and correlated sampling for global sensitivity analysis and generation of pseudomolecule library |
| **Synthetic pseudomolecule library** | This study | N = $10^4$ pseudomolecules | Drug-specific inputs for uncertainty propagation and model agreement analyses |

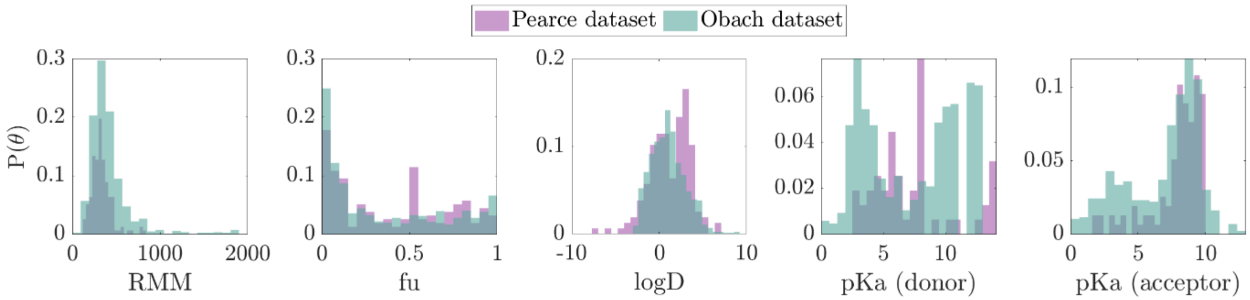


**Figure S1. Drug property distributions in the datasets used for model evaluation**

**Table S2.** Comparative performance metrics for rat *Kp* prediction across 157 compounds.

| Performance metric | all (157 drugs, N=992) | | | | acids (35 drugs, N=258 ) | | | | bases (89 drugs, N=584) | | | |
|---|---|---|---|---|---|---|---|---|---|---|---|---|
| | This model | Mathew | Pearce, calibrated | Pearce, uncalibrated | This model | Mathew | Pearce, calibrated | Pearce, uncalibrated | This model | Mathew | Pearce, calibrated | Pearce, uncalibrated |
| **within 2-fold [%]** | 49 | 36 | 53 | 44 | 57 | 46 | 52 | 46 | 43 | 35 | 53 | 42 |
| **within 3-fold [%]** | 66 | 56 | 72 | 65 | 74 | 66 | 72 | 68 | 60 | 54 | 69 | 63 |
| **within 10-fold [%]** | 92 | 87 | 95 | 91 | 92 | 91 | 96 | 89 | 91 | 86 | 93 | 90 |
| **RMSLE** | 0.40 | 0.48 | 0.34 | 0.42 | 0.26 | 0.27 | 0.23 | 0.34 | 0.48 | 0.57 | 0.40 | 0.48 |
| **AFE** | 0.67 | 0.64 | 0.96 | 1.1 | 0.83 | 0.50 | 1.5 | 1.2 | 0.60 | 0.83 | 0.78 | 1.0 |
| **CCC** | 0.58 | 0.61 | 0.66 | 0.60 | 0.59 | 0.50 | 0.58 | 0.55 | 0.40 | 0.50 | 0.57 | 0.45 |

| Performance metric | neutrals (15 drugs, N=58) | | | | zwitterions (18 drugs, N=92) | | | |
|---|---|---|---|---|---|---|---|---|
| | This model | Mathew | Pearce, calibrated | Pearce, uncalibrated | This model | Mathew | Pearce, calibrated | Pearce, uncalibrated |
| **within 2-fold [%]** | 57 | 40 | 52 | 43 | 56 | 17 | 63 | 52 |
| **within 3-fold [%]** | 72 | 64 | 74 | 67 | 76 | 37 | 84 | 72 |
| **within 10-fold [%]** | 98 | 95 | 97 | 97 | 100 | 80 | 100 | 99 |
| **RMSLE** | 0.29 | 0.30 | 0.26 | 0.42 | 0.24 | 0.38 | 0.21 | 0.27 |
| **AFE** | 0.96 | 0.57 | 1.2 | 1.4 | 0.57 | 0.23 | 0.88 | 0.70 |
| **CCC** | 0.55 | 0.50 | 0.53 | 0.39 | 0.69 | 0.34 | 0.74 | 0.60 |

**Table S3.** Comparative performance metrics for human *Vss* prediction across 862 compounds

| Performance metric | all (N=862) | | | | acids (N=215) | | | | bases (N=309) | | | |
|---|---|---|---|---|---|---|---|---|---|---|---|---|
| | This model | Mathew | Pearce, calibrated | Pearce, uncalibrated | This model | Mathew | Pearce, calibrated | Pearce, uncalibrated | This model | Mathew | Pearce, calibrated | Pearce, uncalibrated |
| within 2-fold [%] | 53 | 47 | 46 | 34 | 65 | 65 | 39 | 32 | 50 | 41 | 50 | 32 |
| within 3-fold [%] | 68 | 65 | 65 | 51 | 77 | 77 | 63 | 52 | 67 | 60 | 67 | 50 |
| within 10-fold [%] | 93 | 90 | 94 | 88 | 96 | 95 | 94 | 88 | 94 | 89 | 94 | 89 |
| RMSLE | 0.32 | 0.40 | 0.32 | 0.42 | 0.21 | 0.22 | 0.22 | 0.37 | 0.39 | 0.49 | 0.41 | 0.50 |
| AFE | 1.2 | 0.95 | 1.3 | 2.4 | 1.2 | 0.84 | 2.0 | 2.8 | 0.97 | 1.0 | 0.82 | 2.2 |
| CCC | 0.58 | 0.59 | 0.47 | 0.51 | 0.58 | 0.58 | 0.37 | 0.44 | 0.36 | 0.50 | 0.24 | 0.25 |

| Performance metric | neutrals (N=227) | | | | zwitterions (N=111) | | | |
|---|---|---|---|---|---|---|---|---|
| | This model | Mathew | Pearce, calibrated | Pearce, uncalibrated | This model | Mathew | Pearce, calibrated | Pearce, uncalibrated |
| within 2-fold [%] | 47 | 47 | 38 | 24 | 49 | 43 | 47 | 36 |
| within 3-fold [%] | 68 | 65 | 65 | 44 | 65 | 62 | 65 | 51 |
| within 10-fold [%] | 97 | 94 | 100 | 79 | 90 | 87 | 94 | 87 |
| RMSLE | 0.24 | 0.25 | 0.24 | 0.44 | 0.32 | 0.43 | 0.29 | 0.40 |
| AFE | 1.3 | 0.77 | 1.6 | 3.2 | 1.3 | 1.0 | 1.4 | 2.2 |
| CCC | 0.38 | 0.41 | 0.25 | 0.25 | 0.46 | 0.45 | 0.43 | 0.46 |

N: number of molecules. AFE: average fold error. CCC: concordance correlation coefficient. RMSLE: root mean squared logarithmic error.

**Table S4.** Prediction variance of *Vss*

| Model | $\sigma^2(Vss)$ [$L^2/kg^2$] - median [95% interval] |
|---|---|
| **This model** | 0.37 [0.35, 0.39] |
| **Mathew** | 0.41 [0.36, 0.47] |
| **Pearce, calibrated** | 0.16 [0.16, 0.17] |
| **Pearce, uncalibrated** | 3.12 [ 2.86, 3.39] |

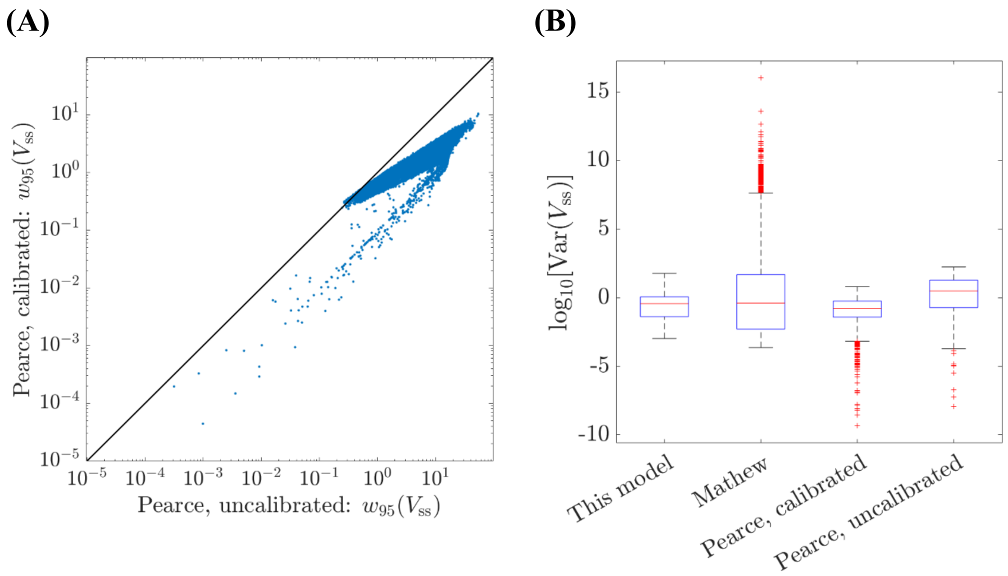


**Figure S2. Model-specific variability in predicted *Vss*.** (A) Impact of calibration on 95% prediction intervals of the Pearce model. Each point represents one of $10^4$ pseudomolecules. Axes reflect the central 95% Monte Carlo prediction interval for the uncalibrated (x-axis) and calibrated (y-axis) forms of the model. The identity line indicates equivalence, with points below the line indicating that calibration reduces the spread of model predictions. Predictions reflect 1000 Monte Carlo realizations per molecule, simulated under identical parameter uncertainty for the calibrated and uncalibrated Pearce models. (B) Variance of predicted *Vss*. Boxplots reflect 1000 Monte Carlo realizations per molecule, simulated under identical parameter uncertainty for each PBPK model. Markers indicate outliers – identified as values lying beyond 1.5 times the interquartile range.

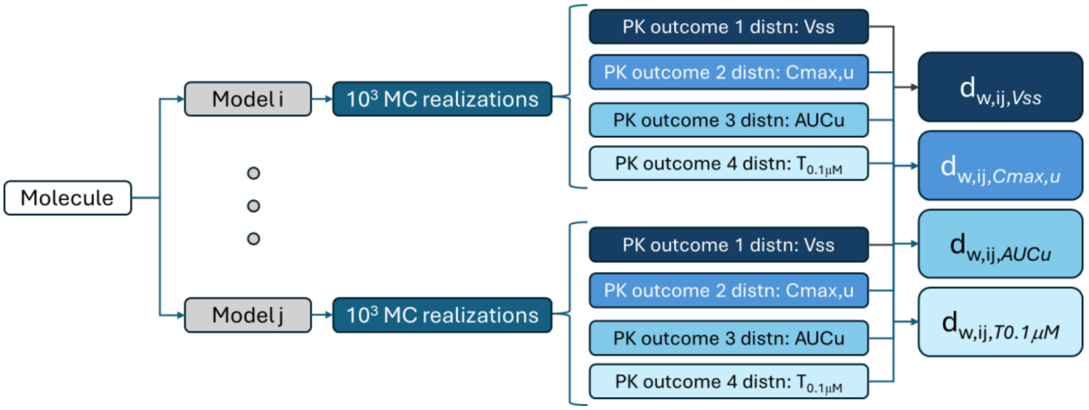


**Figure S3**. The model agreement analysis workflow represents each molecule by Wasserstein distances computed between the Monte Carlo distributions of four PK outcomes for each pair of PBPK models. For the four models considered in this work, that leads to six pairwise comparisons. Thus, the analysis generated 24 Wasserstein distance features for each molecule. We then transformed these features via PCA transformation to group the $10^4$ pseudomolecules in our dataset by K-means clustering of the corresponding $10^4$ twenty-four-element vectors. A similar workflow mapped PBPK predictions for Obach molecules to the 862 twenty-four-element vectors needed to assign Obach molecules to the feature space defined by the pseudomolecule dataset.

**Table S5. K-means sensitivity analyses**

| | | | raw-PCA | raw-Zscore | raw-min-max | whitened-PCA | whitened-Zscore | whitened-min-max |
|---|---|---|---|---|---|---|---|---|
| ADJUSTED RAND INDICES | raw-PCA | | | 0.82 | 0.65 | 0.91 | 0.72 | 0.53 |
| | raw-Zscore | | | | 0.74 | 0.80 | 0.85 | 0.67 |
| | raw-min-max | | | | | 0.64 | 0.77 | 0.84 |
| | whitened-PCA | | | | | | 0.72 | 0.53 |
| | whitened-Zscore | | | | | | | 0.75 |
| | whitened-min-max | | | | | | | |
| | cluster sizes | | 764, 9236 | 1000, 10, 8990 | 1322, 8, 8670 | **751, 9249** | 1175, 2, 8823 | 1609, 12, 8379 |
| SILHOUETTE STATISTICS | minimum | | -0.41 | -0.61 | -0.68 | **-0.42** | -0.95 | -0.52 |
| | quantile | 2.5% | 0.34 | -0.36 | -0.41 | **0.30** | -0.55 | -0.36 |
| | | 50% | 0.94 | 0.76 | 0.68 | **0.94** | 0.82 | 0.62 |
| | | 97.5% | 0.96 | 0.83 | 0.78 | **0.95** | 0.87 | 0.75 |
| | maximum | | 0.96 | 0.84 | 0.80 | **0.96** | 0.89 | 0.76 |
| | mean | | 0.88 | 0.63 | 0.55 | **0.88** | 0.66 | 0.48 |

*Column with bold values corresponds to the selected pre-processing and transformation strategy.

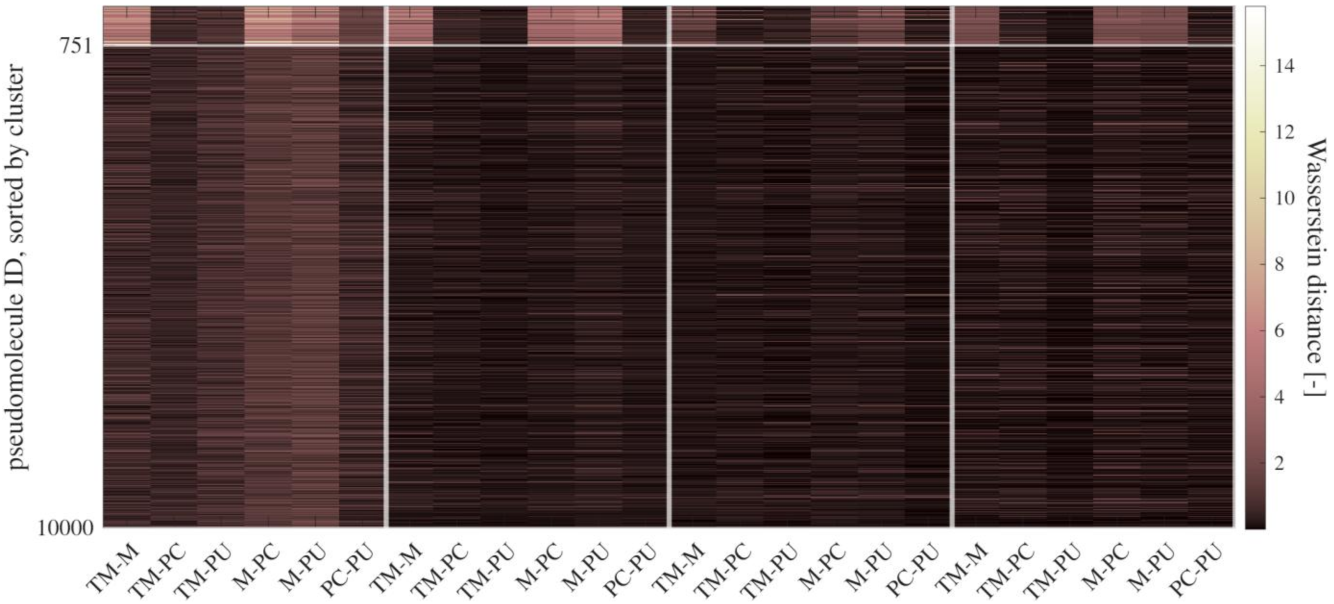


**Figure S4. Heat map of Wasserstein distances between predictions made under uncertainty for specified pairs of models.** K-means clustering segregated$10^4$ pseudomolecules into two groups (Cluster 1: $N_1$=751; Cluster 2: $N_2$=9249). The heatmap shows pairwise Wasserstein distances between model predictions for four outcomes, with columns denoting model pairs and rows indicating individual pseudomolecules, sorted by cluster.

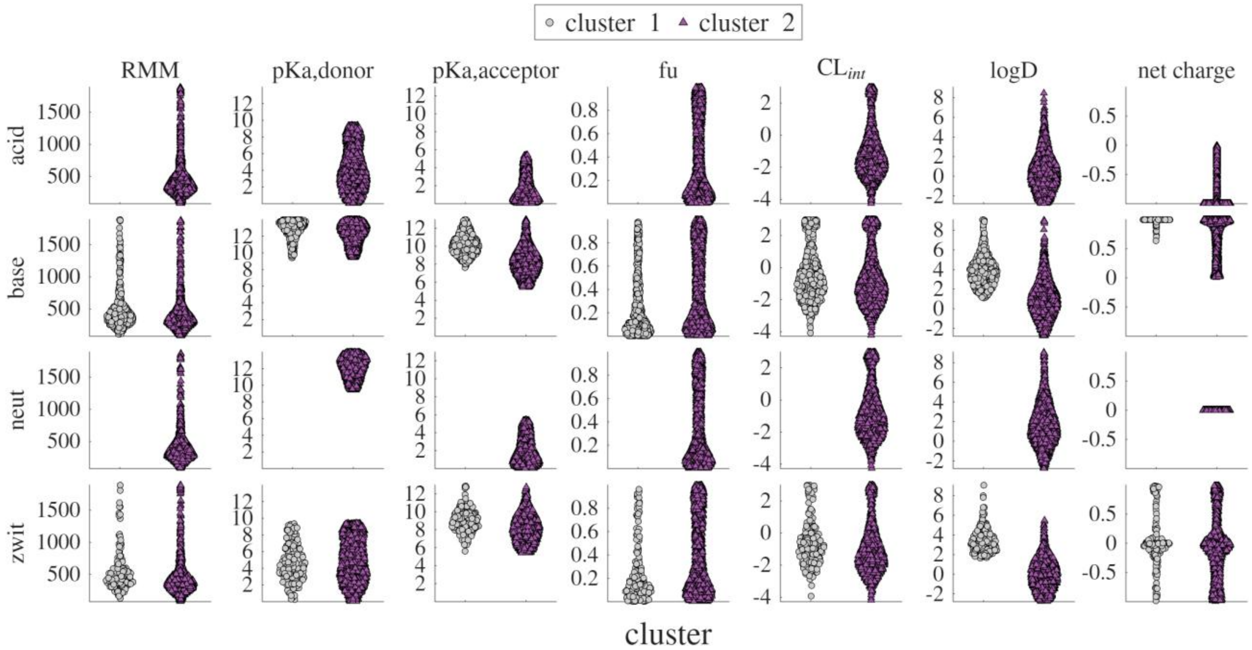


**Figure S5. Physicochemical properties of $10^4$ pseudomolecules grouped by ionization class and cluster.** Each panel shows the distribution of molecular properties within the acid, base, neutral, and zwitterion sub-groups of Cluster 1 ($N_1$ = 751), Cluster 2 ($N_2$ = 9249).

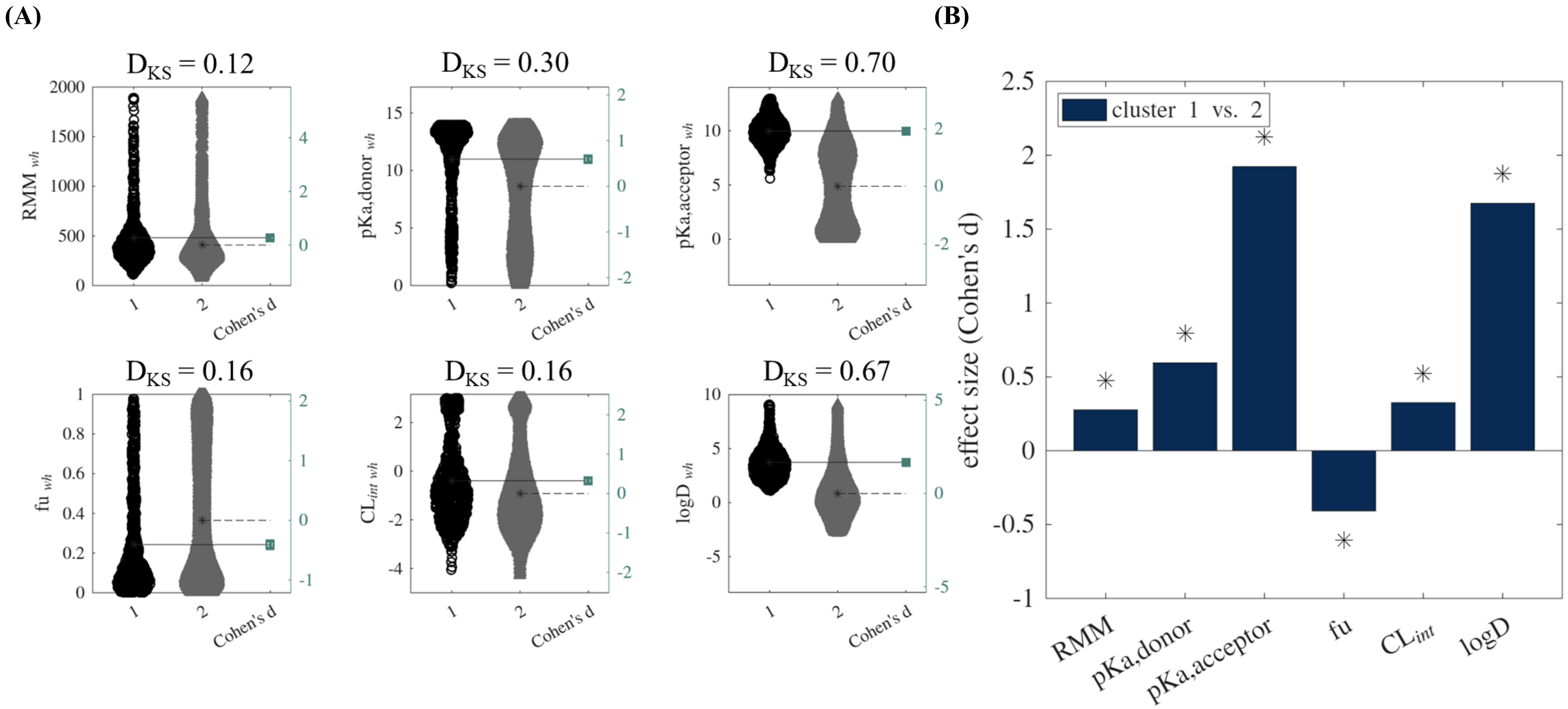


**Figure S6. Effect sizes characterizing the biophysicochemical property differences between pseudomolecules assigned to Clusters 1 and 2.** **(A)** Gardner-Altman plots of properties for molecules in each cluster. Second y-axis indicates the effect size, calculated as Cohen's d values, with bootstrapped confidence intervals. (B) Summary of the Cohen's d effect sizes across all six properties. Asterisks indicate those features with a statistically significant difference between the two clusters (as determined by Kolmogorov-Smirnov statistic, $D_{KS}$, exceeding the critical value $D_{critical} = 0.063$). Number of molecules per cluster: $N_1 = 751$, $N_2 = 9249$.

(A)

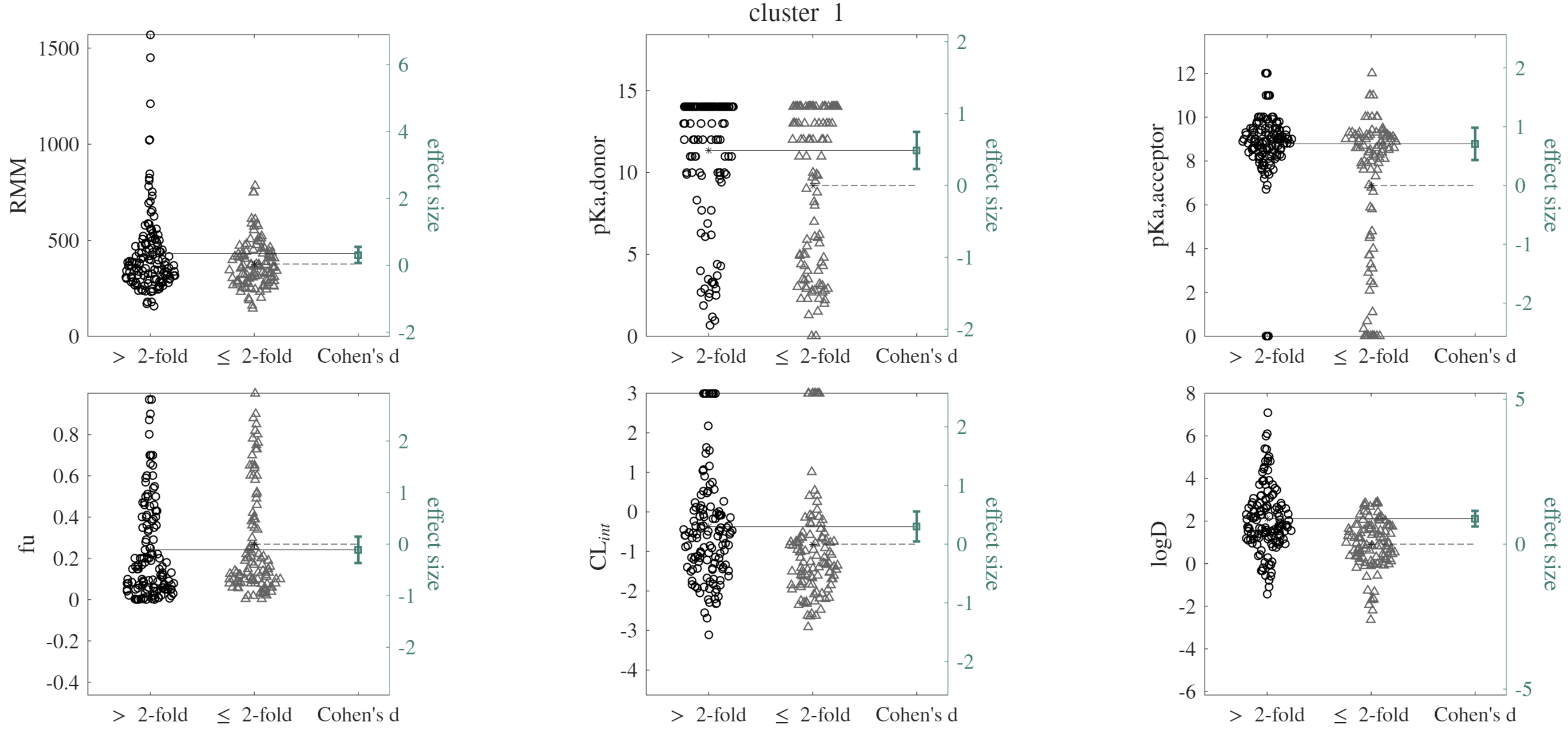
cluster 1
RMM
effect size
1500
1000
500
0
6
4
2
0
-2
> 2-fold
≤ 2-fold
Cohen's d
pKa,donor
15
10
5
0
2
1
0
-1
-2
pKa,acceptor
12
10
8
6
4
2
0
2
1
0
-1
-2
fu
0.8
0.6
0.4
0.2
0
-0.2
-0.4
2
1
0
-1
-2
CL$_{int}$
3
2
1
0
-1
-2
-3
-4
2
1
0
-1
-2
logD
8
6
4
2
0
-2
-4
-6
5
0
-5

**(B)**

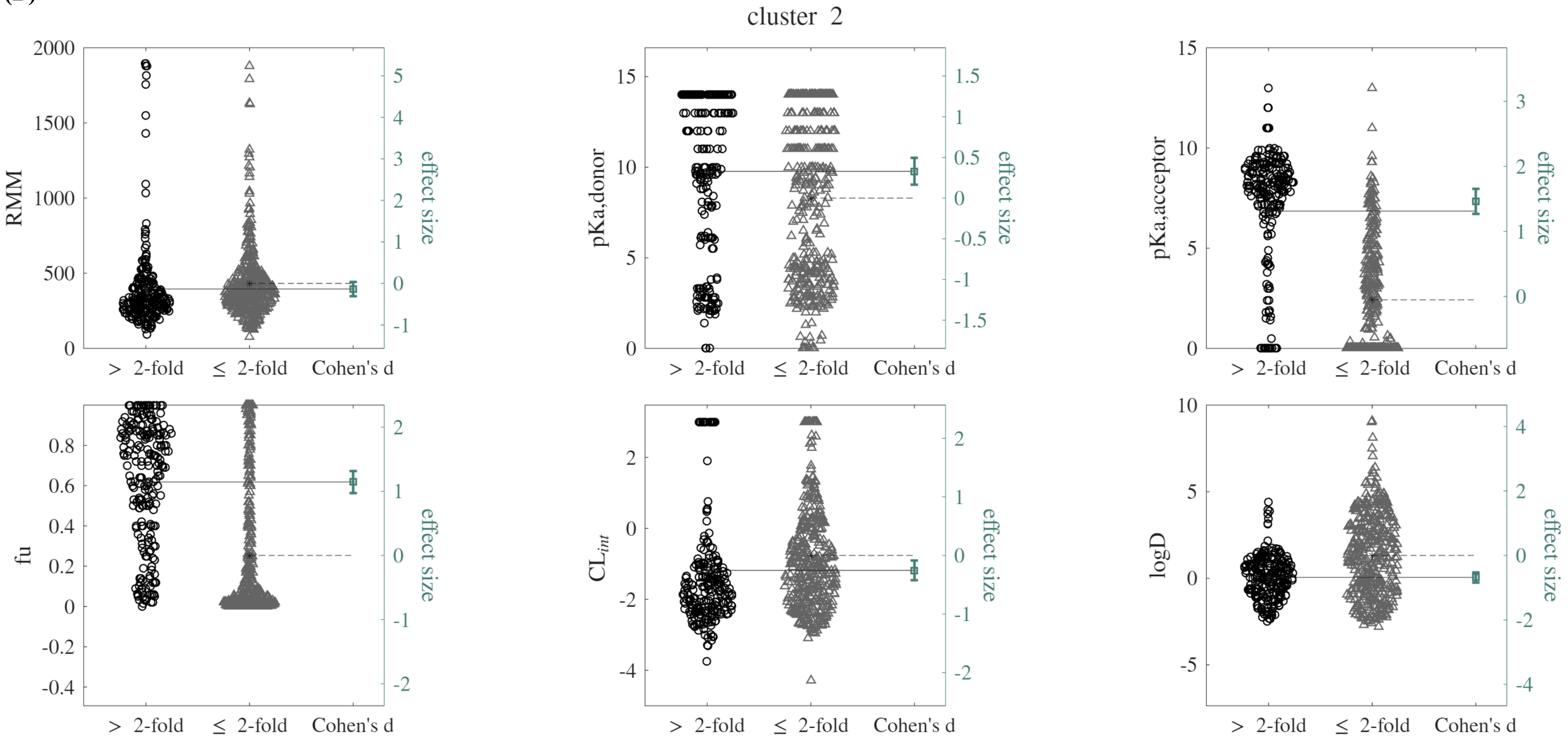


**Figure S7. Effect sizes characterizing the property differences between concordant and discordant prediction subgroups of the Obach molecule clusters.** (A) Cluster 1. (B) Cluster 2. Gardner-Altman plots of properties for molecules in each sub-group. Sub-groups represent molecules for which our model and the Mathew model make nominal *Vss* predictions that differ by greater than two-fold (left) or less than two-fold (right). The second y-axis indicates the effect size, calculated as Cohen's d values, with bootstrapped confidence intervals. **Table S6** reports the corresponding KS statistics and critical values. Number of molecules per cluster: $N_1 = 240$, $N_2 = 622$.

**Table S6. Results from statistical comparison of Obach molecule cluster and subgroup properties.**

| cluster | critical KS value | feature | $D_{KS}$ | effect size (Cohen's d) | cluster | subgroup sizes | | critical KS value | feature | $D_{KS}$ | effect size (Cohen's d) |
|---|---|---|---|---|---|---|---|---|---|---|---|
| **1** | 0.13 | *RMM* | 0.10 | -0.053 | **1** | ≤ 2-fold difference | 108 | 0.21 | *RMM* | 0.12 | 0.31 |
| | | | | | | | | | donor *pKa* | 0.24 | 0.49 |
| | | donor *pKa* | 0.20 | 0.35 | | | | | acceptor *pKa* | 0.28 | 0.71 |
| | | | | | | 2-fold difference | 132 | | *fu* | 0.14 | -0.11 |
| | | acceptor *pKa* | 0.56 | 1.2 | | | | | log $CL_{int}$ | 0.26 | 0.30 |
| | | | | | | | | | *logD* | 0.40 | 0.88 |
| **2** | | *fu* | 0.22 | -0.42 | **2** | ≤ 2-fold difference | 397 | 0.14 | *RMM* | 0.22 | –0.13 |
| | | | | | | | | | donor *pKa* | 0.18 | 0.33 |
| | | log $CL_{int}$ | 0.19 | 0.22 | | | | | acceptor *pKa* | 0.63 | 1.5 |
| | | | | | | 2-fold difference | 225 | | *fu* | 0.51 | 1.1 |
| | | *logD* | 0.29 | 0.39 | | | | | log $CL_{int}$ | 0.25 | –0.26 |
| | | | | | | | | | *logD* | 0.40 | –0.68 |

*KS: Kolmogorov-Smirnov. $D_{KS}$: KS test statistic.

**Table S7.** Biophysicochemical properties of drugs with largest *Kpu* changes due to Pearce calibrations

| | DHA Paclitaxel | Valspodar | Cyclosporine | Anidulafungin | Tirilazad | Ombitasvir |
|---|---|---|---|---|---|---|
| **RMM** | 1164 | 1215 | 1203 | 1140 | 624 | 894 |
| **fu** | 0.0038 | 0.022 | 0.068 | 0.16 | 0.006 | 0.001 |
| **logD** | 9.1 | 9.03 | 8.13 | 7.5 | 7.1 | 7.06 |
| **$pKa_{donor}$** | 12 | 10 | - | 9.2 | - | 13 |
| **$pKa_{acceptor}$** | - | - | - | - | 8.2 | 4.5 |

**Table S8.** Impact of calibration on Pearce model predictions of *Kpu* (quantified as 100 × $Kpu_{calibrated}/Kpu_{uncalibrated}$)

| | DHA Paclitaxel | Valspodar | Cyclosporine | Anidulafungin | Tirilazad | Ombitasvir |
|---|---|---|---|---|---|---|
| **lung** | 134% | 134% | 134% | 134% | 134% | 134% |
| **gut** | 91% | 91% | 91% | 91% | 91% | 91% |
| **spleen** | 89% | 89% | 89% | 89% | 89% | 90% |
| **heart** | 58% | 58% | 58% | 58% | 58% | 59% |
| **muscle** | 82% | 82% | 82% | 82% | 82% | 82% |
| **adipose** | 7% | 7% | 7% | 7% | 7% | 7% |
| **kidney** | 170% | 170% | 170% | 170% | 170% | 171% |
| **skin** | 38% | 38% | 38% | 38% | 38% | 38% |
| **brain** | 31% | 31% | 31% | 31% | 31% | 31% |
| **bone** | 92% | 92% | 92% | 92% | 92% | 92% |
| **liver** | 142% | 142% | 142% | 142% | 142% | 142% |
| **rest of body** | 11% | 11% | 11% | 11% | 11% | 11% |

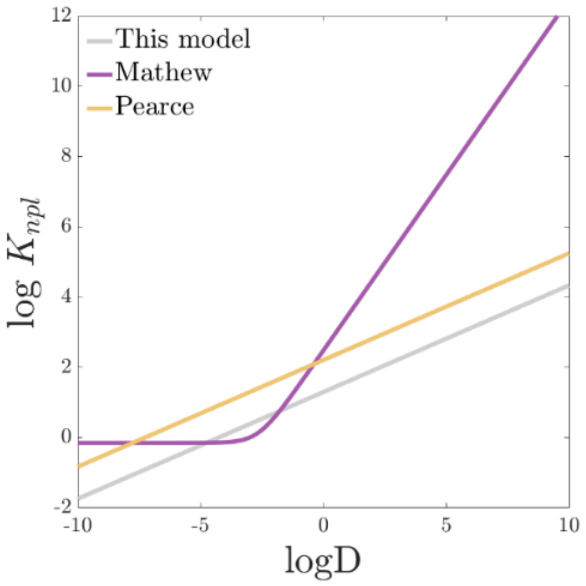


**Figure S8.** Relationship between the neutral phospholipid partition coefficient (*log* $K_{npl}$) and *logD*, as per the assumptions of each PBPK model.

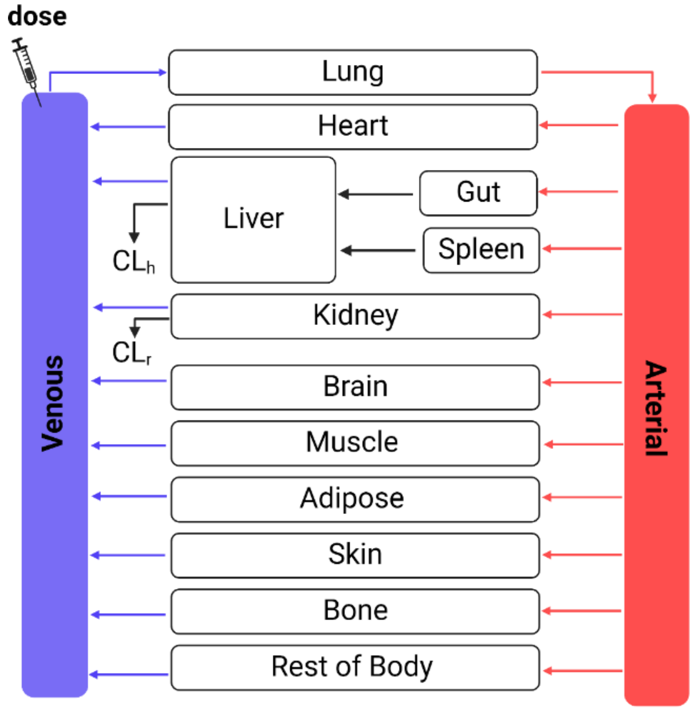


**Figure S9. Schematic of the generic whole-body PBPK model.** Arterial and venous blood circulation connect the major tissues, with intravenous dosing and clearance via the liver and kidney

**(A) This Model, Cluster 1**

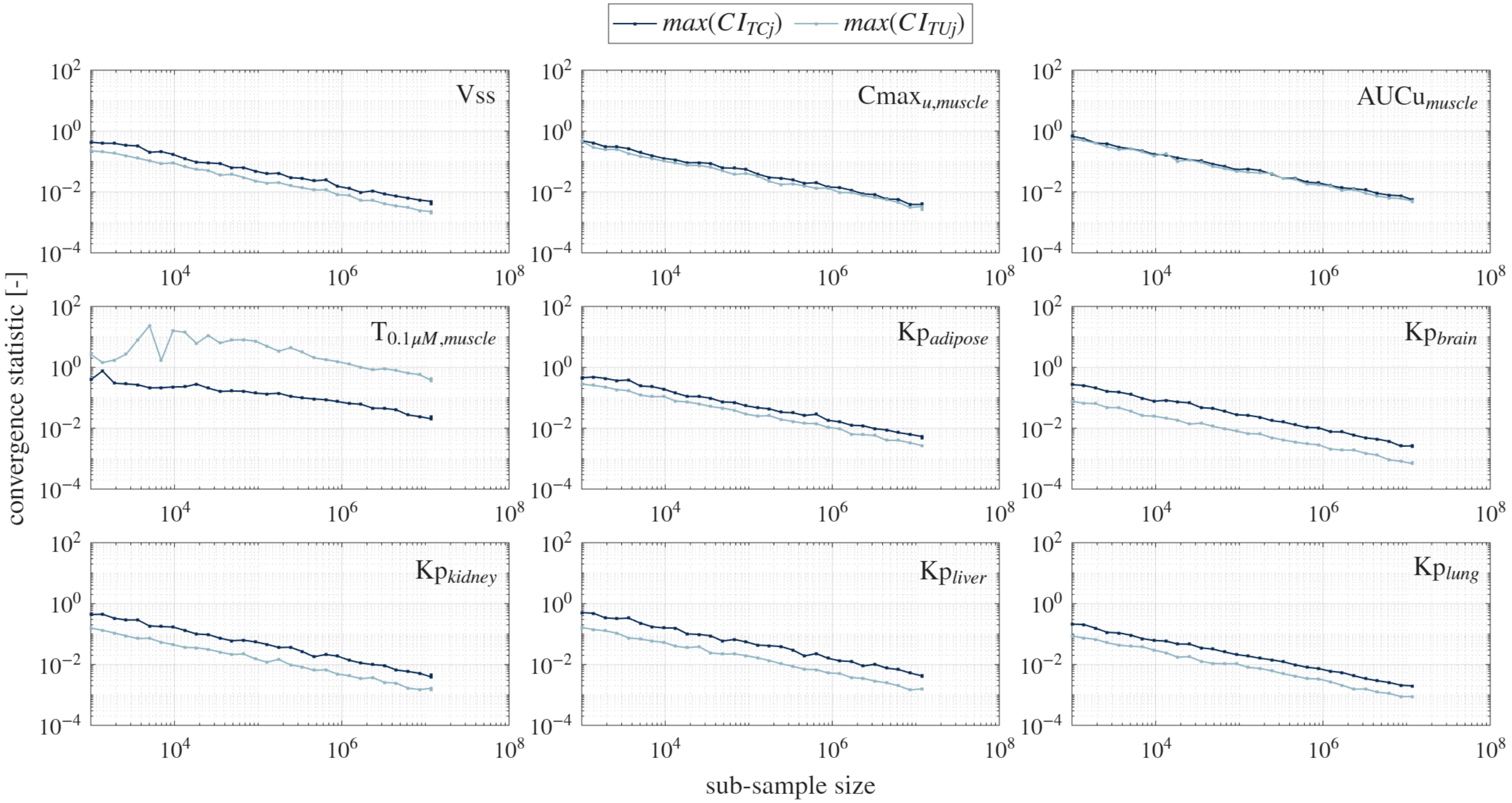

**(B) Mathew Model, Cluster 1**

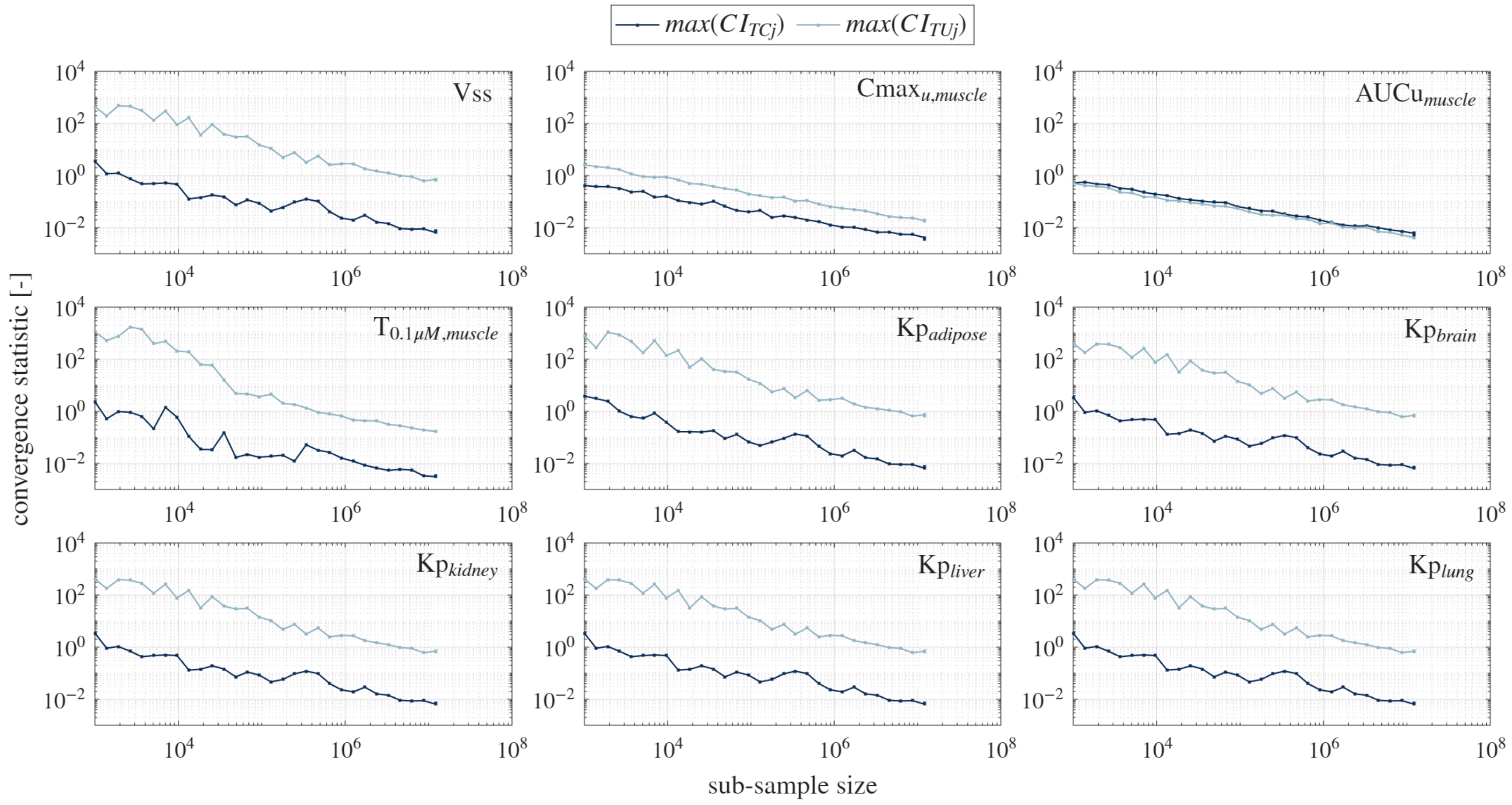

**(C) Calibrated Pearce Model, Cluster 1**

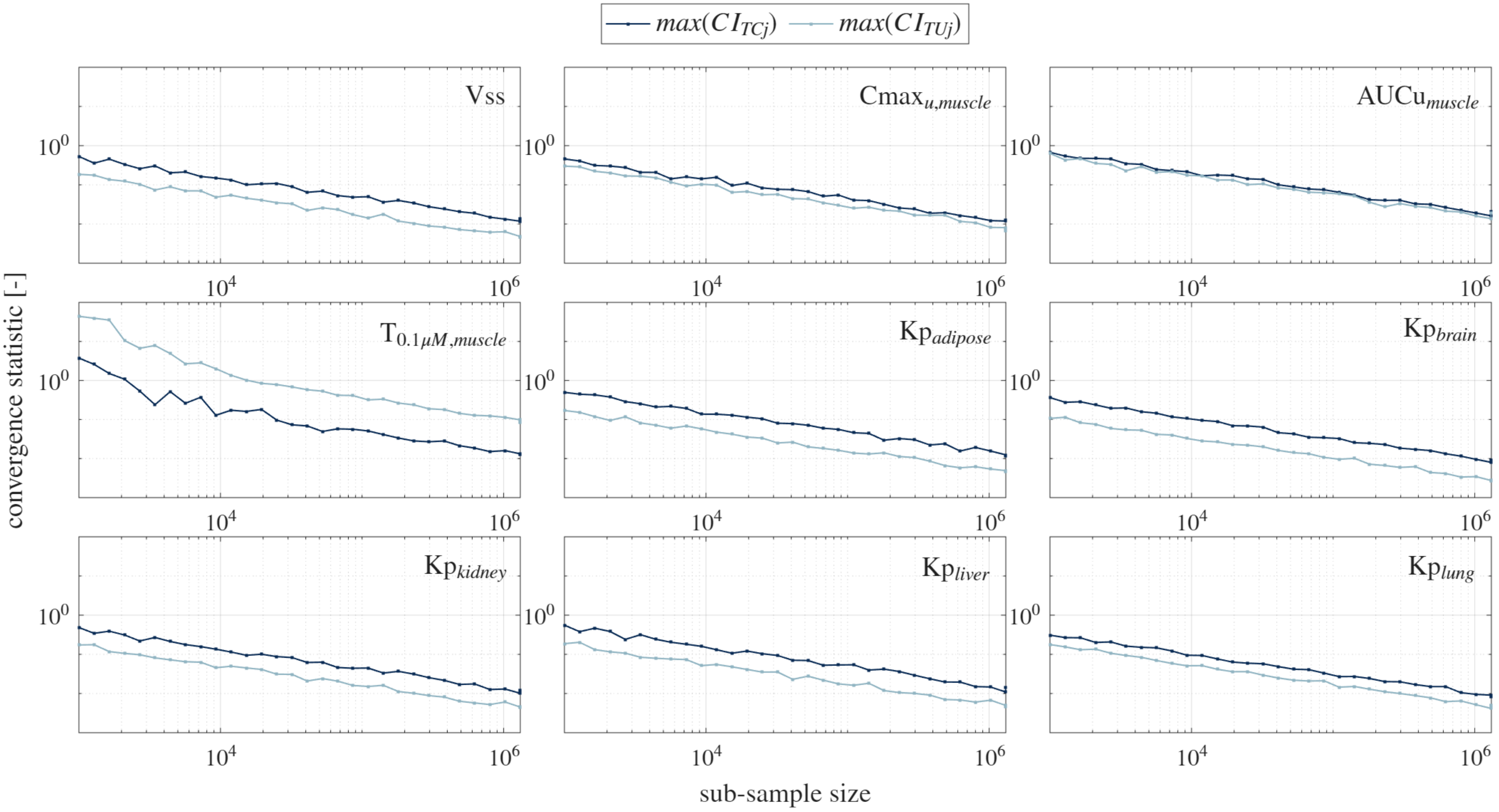

**(D) Uncalibrated Pearce Model, Cluster 1**

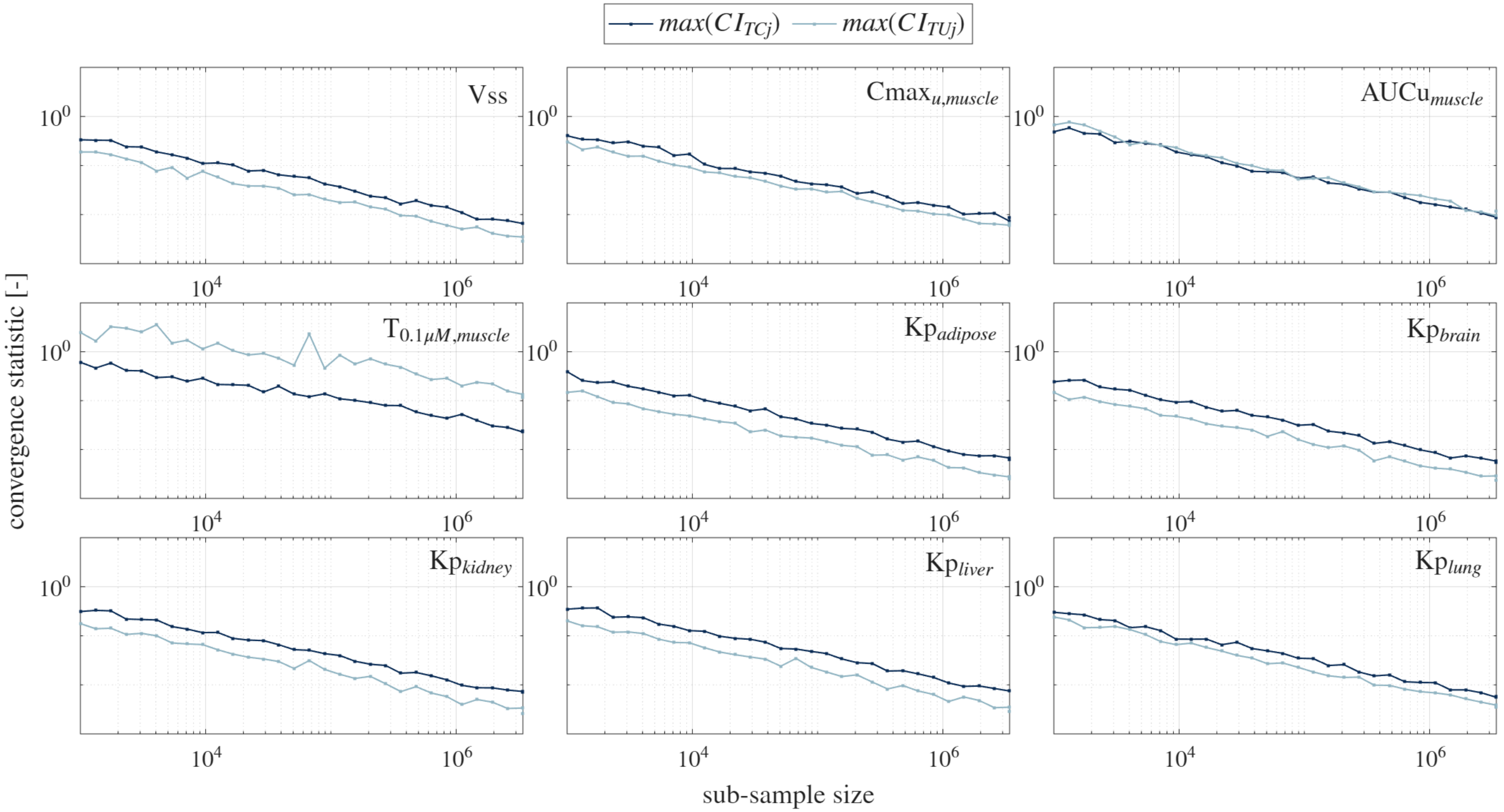


**Figure S10. Sobol' convergence analysis.** Maximum confidence interval for the Sobol' indices of the indicated PK outcome, reported as a function of the number of samples used in the Monte Carlo estimator. Results for Cluster 1 global sensitivity analysis. (A) Our model. (B) Mathew (C) Calibrated Pearce. (D) Uncalibrated Pearce.

**(A) This Model, Cluster 2**

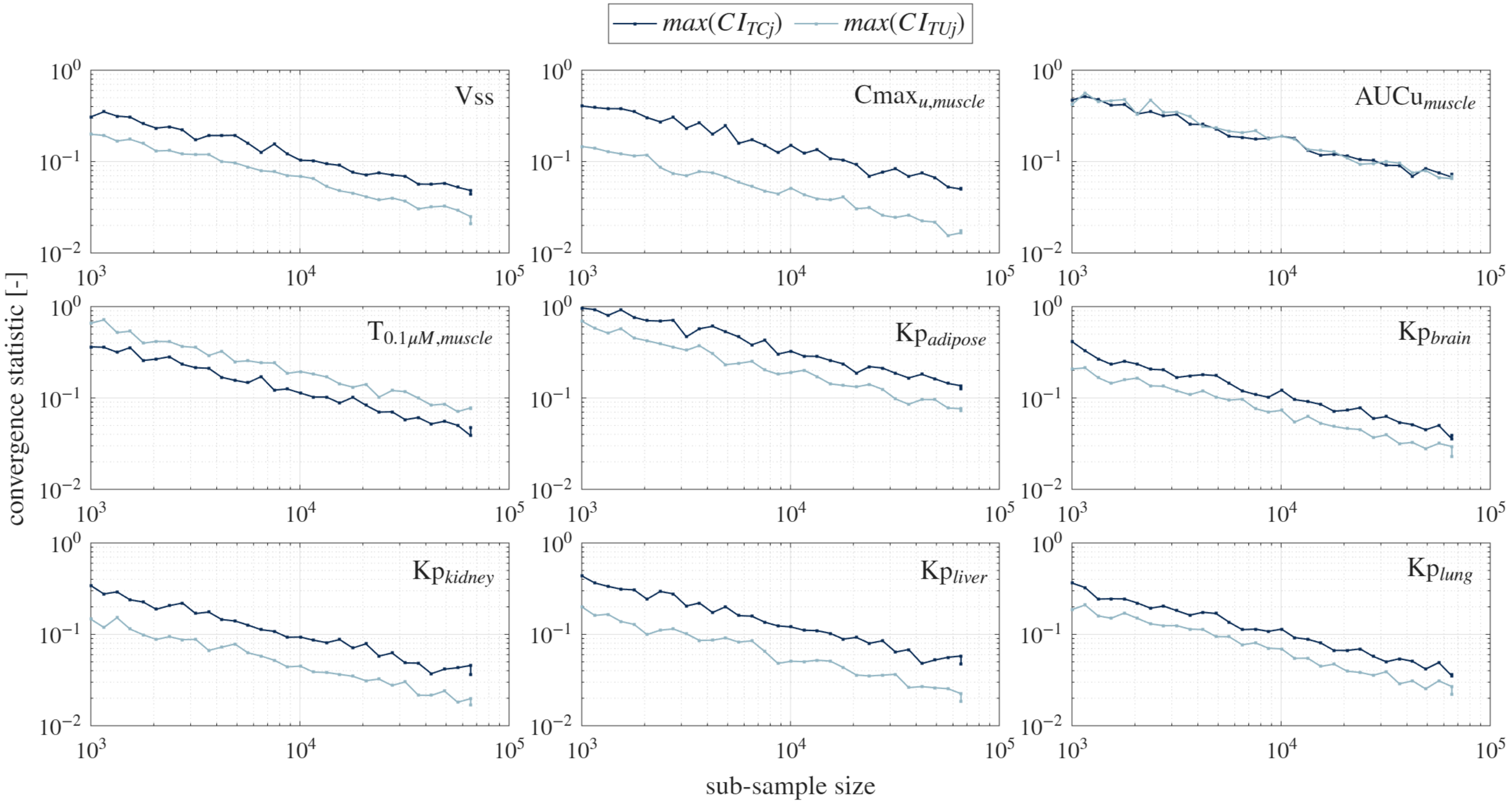

**(B) Mathew Model, Cluster 2**

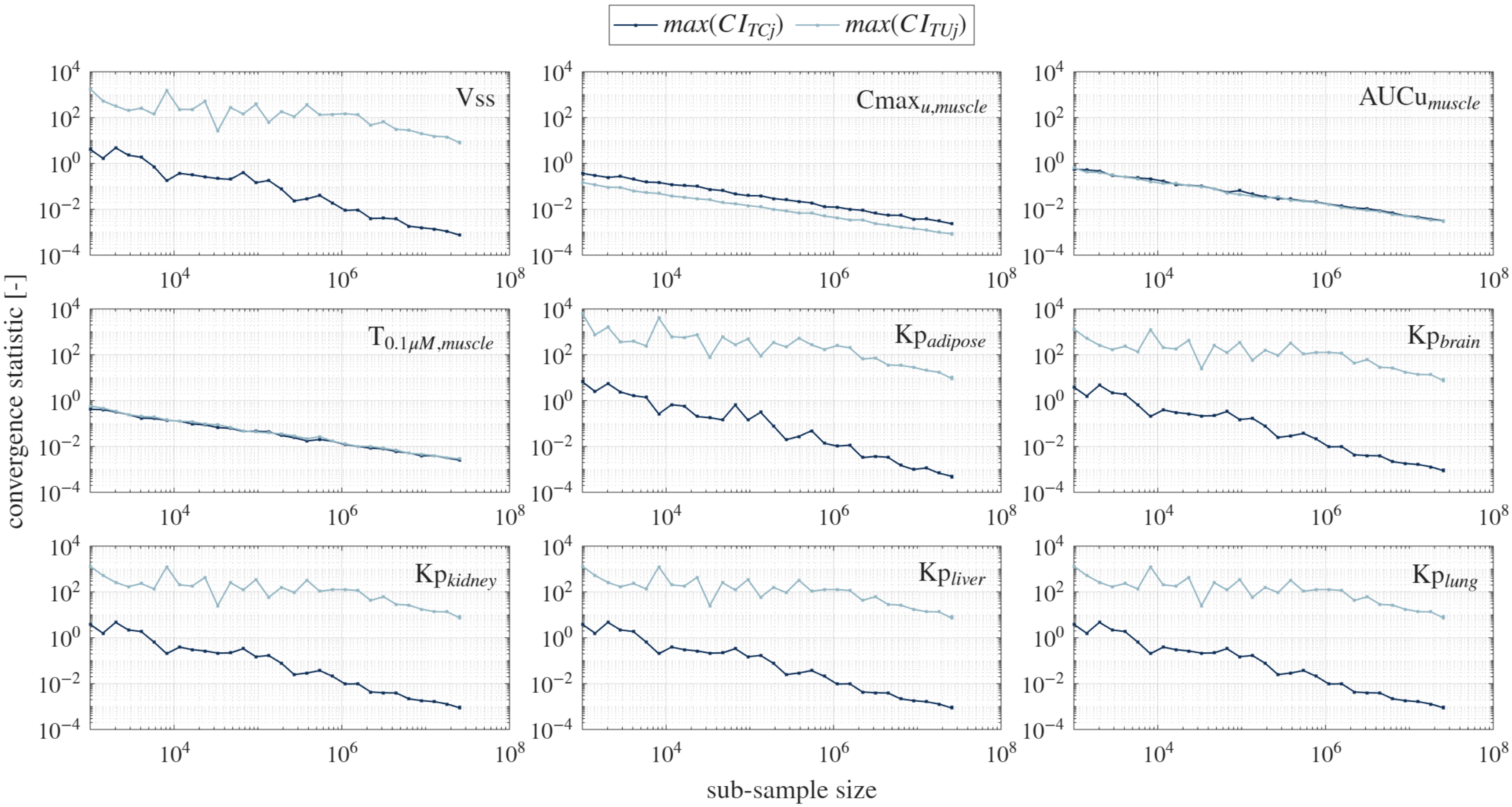

**(C) Calibrated Pearce Model, Cluster 2**

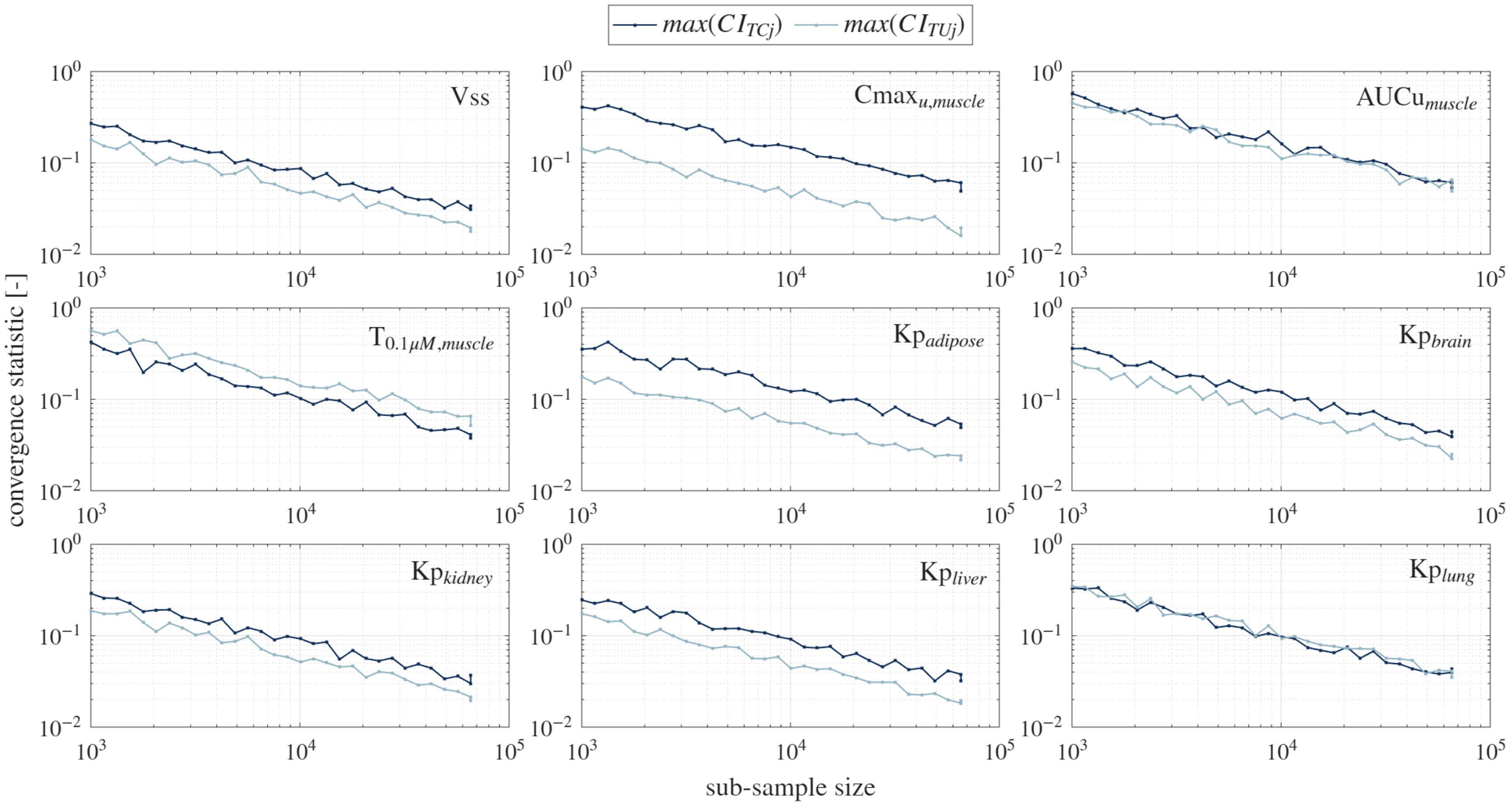

**(D) Uncalibrated Pearce Model, Cluster 2**

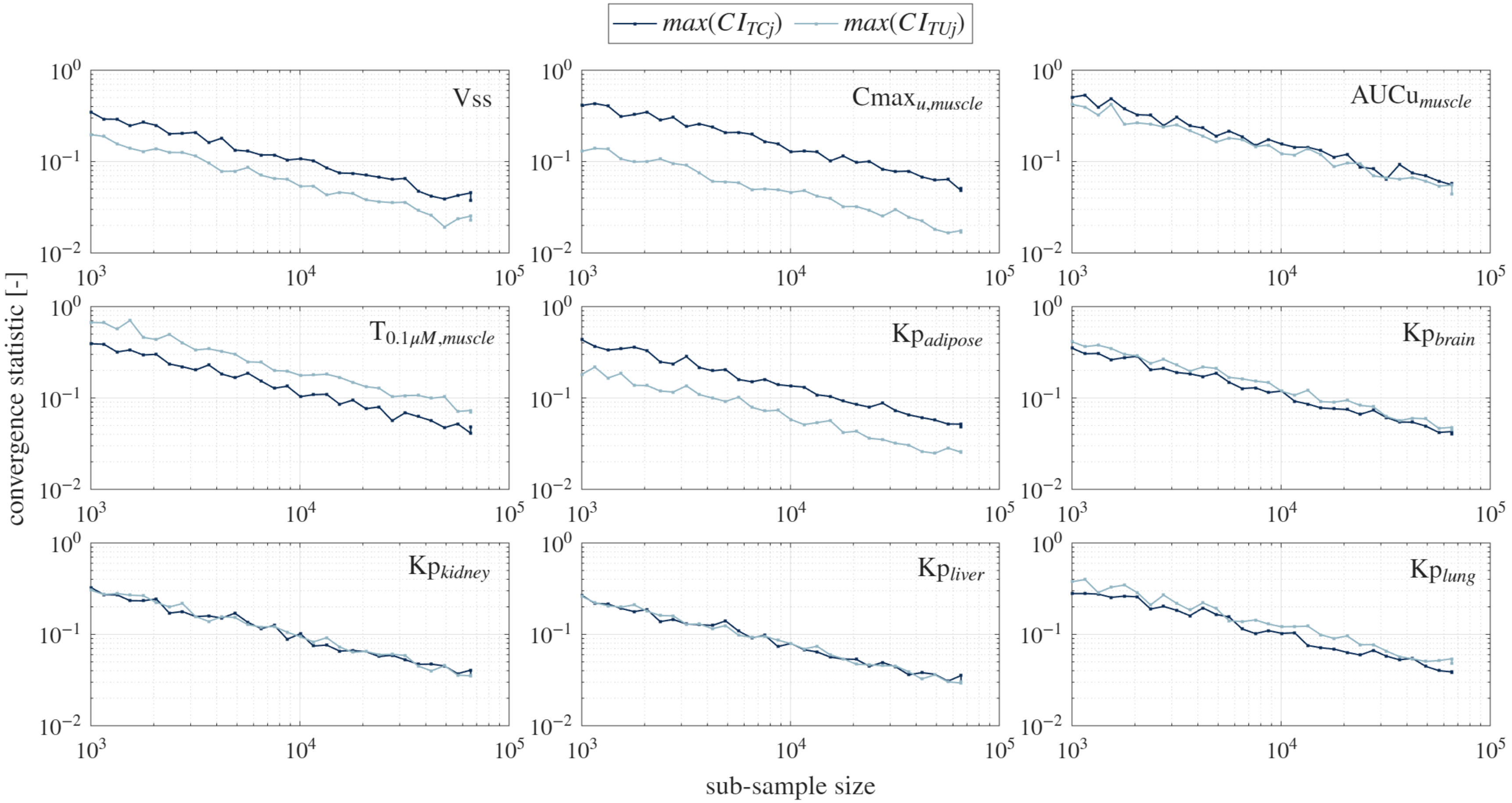


**Figure S11. Sobol' convergence analysis.** Maximum confidence interval for the Sobol' indices of the indicated PK outcome, reported as a function of the number of samples used in the Monte Carlo estimator. Results for Cluster 2 global sensitivity analysis. (A) Our model. (B) Mathew (C) Calibrated Pearce. (D) Uncalibrated Pearce.

**Table S9. Random number generator seeds.**

| Monte Carlo simulations | K-means clustering |
|---|---|
| 52897696 | 1000 seeds (one per K-means clustering attempt) provided as file *rngSeeds.mat*.<br>See code repository at https://github.com/MADBioLab/PBPKPredictionVariability. |
